# Upwelling periodically disturbs the ecological assembly of microbial communities in Lake Ontario

**DOI:** 10.1101/2025.01.17.633667

**Authors:** Augustus Pendleton, Mathew Wells, Marian L. Schmidt

**Author notes:** **Corresponding Author:** Marian L. Schmidt. **Author Contribution Statement:** AP contributed to conceptualization, methodology, software, formal analysis, investigation, visualization, data curation, and writing. MW contributed to methodology, data curation, and writing. MLS contributed to the conceptualization, methodology, investigation, visualization, resources, data curation, writing, supervision, project administration, and funding acquisition. **Study Funding:** Funding for this work was made possible via a grant to MLS from the Affinito-Stewart & President’s Council of Cornell Women, Cornell University start-up funds to MLS, and a NOAA Margaret A Davidson Fellowship to AP (NA24NOSX420C0016). **Data Availability Statement:** All raw and processed data for this project are publicly available. The main GitHub repository for this project is available at https://github.com/MarschmiLab/Pendleton_2025_Ontario_Publication_Repo, which includes all processed data (Amplicon Sequence Variants (ASVs), CTD casts, water quality data, all other metadata, etc.), the code for all figures, tables, and summary statistics, including the generation of ASVs using the DADA2 workflow, and a reproducible *renv* environment. All the software versions and citations used for data processing and statistics are described in Table S2. The raw compressed 16S rRNA gene sequencing fastq files are available in the NCBI Sequence Read Archive under the BioProject accession numbers PRJNA1212049. All flow cytometry data are available in the FlowRepository database with the ID number FR-FCM-Z8SJ.

## Abstract

The Laurentian Great Lakes hold 21% of the world’s surface freshwater and supply drinking water to nearly 40 million people. We provide the first evidence that wind-driven upwelling restructures microbial communities in Lake Ontario, with its effects sustained and redistributed by an internal Kelvin wave propagating along the shoreline. We combine 16S rRNA metabarcoding, absolute abundance quantification via flow cytometry, and hydrodynamic profiling to link physical processes to community composition. While thermal stratification organizes microbial communities by depth and season, this vertical structure arises from contrasting mechanisms: homogenizing selection in surface waters and dispersal limitation and drift in the hypolimnion. Kelvin wave-driven upwelling disrupts this scaffold, displacing rare taxa into the surface and creating novel coastal communities predicted to be enriched in methane oxidation and sulfur metabolism genes—functional traits absent elsewhere in the lake. We observed a Kelvin wave lasting over two weeks and propagating eastward at ∼60 km day⁻¹. Given the ∼10–12 day recurrence of wind events during the stratified season, at any time at least one segment of Lake Ontario’s coastline is experiencing upwelling. These recurrent upwellings, sustained and redistributed by Kelvin waves, remodel microbial communities on ecologically relevant timescales. They act as a biological disturbance overriding stratification, mobilizing rare functional potential, and assembling novel coastal microbial communities. As climate change lengthens and intensifies stratified periods and reshapes large-lake circulation, understanding how physical forcing governs microbial assembly is essential for forecasting the biogeochemical future of Earth’s great lakes—especially in shoreline zones where ecological shifts directly affect human communities.

## Introduction

Microbial communities fuel aquatic ecosystems by transforming energy, cycling nutrients, purifying water, and anchoring aquatic food webs [1–4]. In lakes, microbes recycle organic matter and support higher trophic levels via the microbial loop [5]. Yet in Earth’s largest lakes, we still lack a predictive understanding of how microbial communities are structured spatially. Likewise, we do not know how they respond to the powerful physical forces of stratification and circulation that define these systems and are shifting under climate change [6–9].

Classic ecological theory holds that community assembly reflects a balance between deterministic forces like environmental selection and stochastic processes such as drift and dispersal [3, 10]. In hydrodynamically active systems—like Earth’s largest lakes and the global oceans—these ecological processes unfold within a dynamic physical environment shaped by ecosystem-scale drivers like stratification, currents, and wind-driven upwelling [6, 11, 12]. Large lakes, upon which hundreds of millions of people depend, thus offer a tractable but underused model for understanding how physics rewires microbial biogeography and biogeochemical function, with insights that scale to the oceans, where mesoscale circulation shapes the global cycling of carbon, nitrogen, and sulfur [13, 14].

The community assembly mechanisms of selection, dispersal, diversification, and drift interact dynamically in response to environmental heterogeneity [3, 4, 15]. Selection may homogenize communities under stable conditions or drive divergence across gradients, while stochastic processes (e.g. dispersal and drift) introduce variation in community structure unlinked to environmental pressures [10]. These processes shift across spatial and temporal scales, and may respond strongly to physical mixing or isolation. Although microbial communities in smaller lakes and along watershed gradients have been widely studied [16–19], studies of large lakes have more often focused on surface waters or treated them as spatially homogeneous systems [20, 21]. As a result, the influence of vertical structure and hydrodynamic circulation on microbial community assembly in large lakes remains underexplored [22–25]. Planktonie microbes are carried by currents and turbulence, emphasizing the need to understand large lake microbial communities through the dynamics of ecosystem-scale physical processes.

One powerful but largely overlooked driver of microbial dynamics in large lakes is the wind– upwelling–Kelvin wave cascade which occurs throughout the stratified season in large lakes. Strong wind initiates coastal upwelling through Ekman transport, tilting the thermocline. When the wind relaxes, this displaced density interface releases its stored energy as baroclinic internal waves. In offshore waters, these manifest as near-inertial oscillations [26]; along the coast, they generate internal Kelvin waves which are coastally trapped gravity waves that roll anticlockwise in the Northern Hemisphere within a narrow 3–5 km shoreline band [27–29]. Propagating with the wave is a baroclinic coastal jet: a shore-parallel current, strongest near the surface, that redistributes upwelling and downwelling water along the lake’s perimeter, creating a disruption of the thermocline that can last for weeks until the next wind event creates a new upwelling. This process repeats in Lake Ontario every ∼10–12 days [30] and every 5–10 days in Lake Erie [29]. While physical limnologists have long studied these features for their role in transporting nutrients, pollutants, and thermal discharges [11, 27], their ecological consequences remain largely unexamined [31].

Here, we leverage Lake Ontario, a 19,000 km² freshwater basin with well-characterized seasonal circulation, to test how physical and ecological forcings shape microbial community structure and function. The lake stratifies annually and experiences wind-driven upwelling events that transport cold, nutrient-rich water to surface zones [6, 32–34], creating transient and chemically distinct habitats. Using an interdisciplinary framework that integrates V4 16S rRNA metabarcode sequencing, flow cytometry, environmental profiling across 47 stations and two seasons, and physical limnology, we reveal that Kelvin wave–driven upwelling fundamentally reshapes microbial biogeography, mobilizing rare microbes with unique functional potential. We show that microbial communities stratify by depth due to homogenizing selection, not dispersal limitation, but when upwelling disrupts this scaffold, it assembles novel shoreline communities. Together, these findings reveal how physical forcing structures microbial ecosystems and challenge assumptions of spatial homogeneity in large lakes. Crucially, these transformations occur along shorelines that are often densely populated and essential for recreation and drinking water.

## Methods

### Sample collection

Samples were collected from multiple stations throughout Lake Ontario aboard the *R/V Lake Guardian* in May and September of 2023 as part of the US Environmental Protection Agency’s (EPA) Lower Food Web Cooperative Science and Monitoring Initiative (CSMI) survey (Fig. 1A; Table S1). The stations were arranged in 5 north-south transects across the lake, which were sampled in an east-west fashion over May 15^th^-19^th^ and September 23^rd^-27^th^. Water samples were collected using a rosette while depth profiles were taken with a conductivity, temperature, and depth (CTD) meter equipped with fluorometer, dissolved oxygen, spherical photosynthetic active radiation sensor, and turbidity sensors. A total of 30 stations were visited each cruise for CTD casts and EPA-led water chemistry; microbial samples were collected at 15 of these stations. Depending on the depth of a given station, three possible depths were sampled. Surface samples (“E”) were from 5 m below the surface. Mid samples (“M”) were collected at stations which were over >30 m deep, either at the fluorescence maximum or thermocline in May (typically 10 m to 25 m, Table S1), or at the station’s average depth in September. Bottom samples (“B”) were taken 2 m above the lake floor. At every depth sampled, two biological replicates were collected from separate Niskin bottles but reflect the same parcel or water and processed separately. Samples were pre-filtered consecutively through sterilized 200 𝜇m and 20 𝜇m Nitex mesh (Wildco) into sterile 10L bottles (Nalgene). Microbial samples were filtered onto a polyethersulfone 0.22 𝜇m filter (MilliporeSigma). Filters were flash frozen in liquid N_2_ and stored at -80°C.

**Figure 1.**
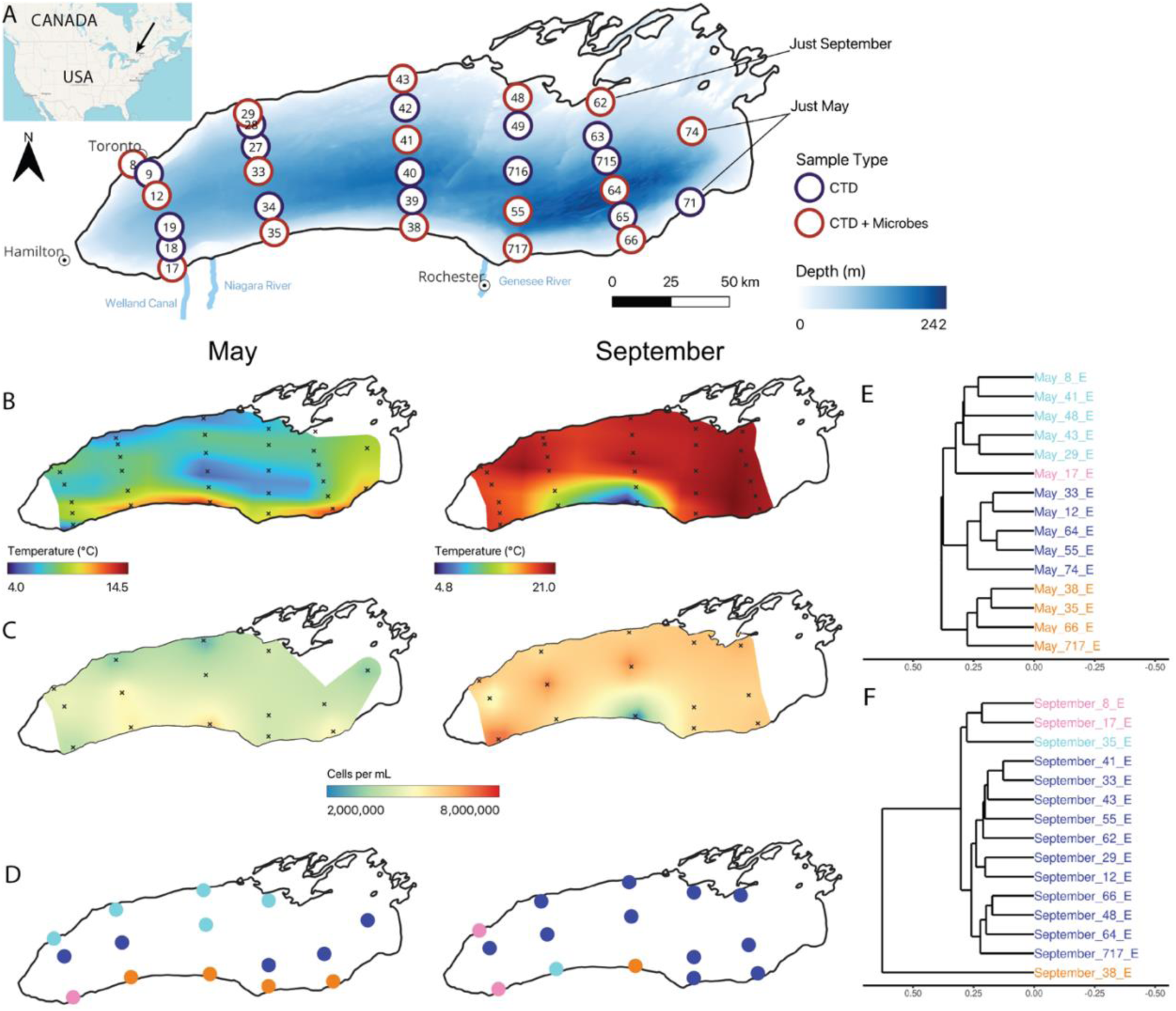
Upwelling reshapes surface microbial abundance and composition. (A) Thirty stations were visited for CTD-casts and EPA-led water chemistry (blue and red), and microbial samples were collected at fifteen stations (red) in both months, though station 62 was only sampled in September, and stations 71 and 74 only in May. CTD data was unreliable for station 65 in September. (B) Surface (5 m) temperatures in Lake Ontario interpolated spatially across the lake in May and September using multilevel B-splines. (C) Surface cell counts interpolated across May and September using multilevel B-splines. (D) Surface community clusters (shown by color) formed via UPGMA clustering on the community dissimilarity (*GU^A^_0.5_*). Trees were cut into four groups for both months, as shown for May in (E) and September in (F). X-axes (tree heights) are consistent between plots.

### Environmental data

Water chemistry and chlorophyll a data were generated by the U.S. EPA’s Great Lakes National Program Office according to their standard protocols [35]. See the Supporting Information for details on environmental data, including temperature monitors and meteorological records.

### Quantifying cells with flow cytometry

Flow cytometry samples were prepared in the field by adding 1 𝜇L (in May) or 5 𝜇L (in September) of 25% glutaraldehyde to 1 mL of sample water with a 10 minute incubation, before flash freezing in liquid nitrogen and later stored at -80 °C. Fixed samples were thawed for 20 minutes at 37 °C, diluted 20X, and stained in triplicate with SYBR I green dye at a concentration of 1x SYBR I green at 37 °C for 20 minutes in the dark. Cell abundance was measured on a BD Accuri C6 Flow Cytometer. Sample volume was set at 50 uL, but actual volume uptake was estimated using CountBright Absolute Counting Beads (Thermofisher) to be 42.5 ± 0.5 𝜇L. Samples were run at a flow rate of 33 uL/min. Fluorescence was measured using the blue laser FL1 filter. Raw events were filtered with a FSC-H filter of 100 and FL1-H filter of 400. Flow data was analyzed using the R packages flowCore and ggcyto [36, 37]. A polygon gate was defined to select for fluorescently-labeled cells relative to unstained controls, allowing some background fluorescence since many environmental cells are naturally pigmented (Fig. S1). This gate is available within the Github repository associated with this manuscript.

### DNA extraction & Illumina sequencing

All DNA extractions were carried out using the Qiagen DNeasy PowerWater kit, mostly following the manufactuer recommendations (more details in Supporting Information). Half a filter was used for May samples; a quarter filter produced sufficient DNA yields and as such was used for September samples, translating to a minimum of 540 mL extracted, maximum of 4380 mL extracted, and a median of 1320 mL extracted. An extraction negative was produced for each kit using a blank filter.

The V4 region of the 16S rRNA gene was amplified using the Earth Microbiome Project protocol [38, 39]. Polymerase chain reaction (PCR) was performed, with blanks, in triplicate 25 𝜇L reactions using KAPA HiFi 2X MasterMix (Roche), 0.2 𝜇M 515F (5’-GTGYCAGCMGCCGCGGTAA) and 806R (5’-GGACTACNVGGGTWTCTAAT) primers with Illumina Nextera adapter overhangs, and 5 ng of sample DNA. A ZymoBIOMICS Microbial Community DNA Standard (Zymo) was included to assess the sequencing error rate. Altogether, 145 environmental samples were sequenced, in addition to two negative field controls to verify contamination from filtration, two extraction negatives, a PCR blank (pooled negative reactions across all PCR plates), the ZymoBiomics positive control, and an indexing negative. Samples and indexing blanks were sequenced using a 2 x 250 bp paired-end strategy on an Illumina MiSeq at the Cornell Biotechnology Resources Center, generating 15.72 million reads with a minimum of 22,430 reads, a maximum of 122,493 reads, and a median sequencing depth of 105,500 reads (excluding blanks).

### Microbial bioinformatic processing

Raw Illumina sequences were processed into amplicon sequence variants (ASVs) using the standard workflow in the dada2 package in R [40]. The phyloseq package was used to organize the ASV count table, metadata, and later taxonomic table and phylogeny [41]. Taxonomy was first assigned using the TaxAss freshwater database [42] and then with the Silva v138.1 16S rRNA database [43]. ASVs classified as mitochondrial and chloroplast (1092 ASVs), and ASVs with a higher relative abundance in any of the negative controls (six ASVs) were removed. In the mock community, three spurious ASVs were detected alongside all nine expected ASVs, potentially arising from sequencing error, index hopping, or contamination during processing. These spurious ASVs represented 0.03% of all reads in the mock community. No environmental samples were discarded based on sequencing depth, leaving 145 samples. The resulting minimum number of reads in a sample was 7,837 reads, a maximum of 72,631 reads, a median of 40,376 reads, and a total of 7,280 unique ASVs.

A phylogenetic tree was constructed using MAFFT for alignment and FastTree under a generalized time-reversible model [44, 45]. The tree was rooted at the most-recent-common-ancestor (MRCA) of the Archaea using the ape package [46]. Several polyphyletic, anomalous tips had exceptionally long branch lengths and no taxonomic assignment and were removed by filtering for tips with node heights greater than 2 as calculated using the phytools::nodeheight function [47].

### Ecological Statistics

Between-sample (beta) diversity was calculated using functions from the vegan or GUniFrac packages (modified to take into account absolute abundances) [48–50]. Unless otherwise noted, community dissimilarity refers to the absolute, generalized weighted Unifrac distance with 𝛼 = 0.5 (*GU^A^_0.5_*, [50]). The relative abundance of each ASV in a given sample was normalized by the cell concentration of that sample, providing the absolute abundance of each ASV. Ordinations were constructed using principal coordinates analysis implemented in phyloseq [41]. Replicates were highly similar to each other (Fig. S2) and were merged by summing reads, resulting in a minimum of 13,986 reads and a median of 82,765 reads. Depth-month groups were defined with UPGMA clustering based on community dissimilarity, cutting the tree at 3 groups (Fig. S3A). Variance partitioning and post-hoc tests were performed with Vegan functions varpart, dbrda, and anova.cca [48]. Permutational multivariate analysis of variance (PERMANOVA) and multivariate homogeneity of groups dispersions tests were run with adonis2 and betadisper from the vegan package [48].

Within-sample (alpha) diversity was estimated using Hill numbers via the iNEXT package [51, 52]. All rarefaction curves were asymptotic, confirming sufficient sequencing depth for representative sample richness (Fig. S4). Differential abundance of microbial ASVs between depth-month groups was calculated at the Class level using the pairwise setting in the ANCOM-BC2 package [53]. Significant differences had an FDR-corrected p-value less than 0.05 and passed the sensitivity test across multiple pseudocount values; other parameters were kept at default settings. Microbial community assembly processes were quantified with iCAMP [54, 55], where *bin.size.limit* specifies the minimum number of ASVs per phylogenetic bin and *ds* defines the initial phylogenetic distance used to define bins. Full details and explanation of iCAMP are available in the supplemental. We used a minimum bin size of 24 ASVs, a maximum bin distance of 0.5, and a confidence threshold of 0.975 compared to a randomized null distribution with 1000 iterations [56]. Results from iCAMP were resilient across ranges of both maximum bin distances (*ds*) and minimum bin sizes (*bin.size.limit*) (Fig. S5). Microbial functional traits were inferred using FAPROTAX [57], using the absolute abundance of each ASV.

When defining “rare” taxa specifically (as in Fig. S10), we refer to them as ASVs with a relative abundance less than 0.01%. Elsewhere in the manuscript, we use unweighted measures of β-diversity, which inherently emphasize differences among rare taxa without applying an explicit abundance threshold.

Statistical testing, including Two-Sample Wilcoxon tests following Kruskal-Wallis tests; correlation tests; and linear models were done in R, using base functions and the ggpubr package [58]. If multiple comparisons occurred within a given plot panel, all p-values were corrected with a Bonferroni Correction.

### Spatial Analysis

Mapping and spatial interpolation was performed either in R using the packages sf, terra, and tmap, or in QGIS using GRASS [59–63]. Data files for Lake Ontario’s bathymetry were downloaded from the National Oceanic and Atmospheric Administration [64].

## Results

### Upwelling events are frequent and extended drivers of spatial heterogeneity in coastal zones

We detected upwelling events of cold water along the northern shore in May (stations 29 & 43) and the southern shore in September (stations 35 and 38) (Fig. 1B, Fig. S6). In May, the warm southern shore likely reflected a combination of downwelling and relatively warmer water flowing in from the Niagara River (Fig. S7A-C). In September, strong easterly winds in the days leading up to sampling likely drove the pronounced southern upwelling observed at station 38, as reflected in both temperature (Fig. 1B, Fig. S7D) and nutrient data (Fig. S8). While prevailing winds in Lake Ontario are typically westerly and result in upwelling events near Toronto, these unique easterlies reversed the usual pattern, generating upwelling near Rochester instead. We estimate this upwelling formed near the Niagara outlet on September 23^rd^ and propagated eastward as a Kelvin wave along the southern shore, concluding at the eastern terminus by October 9^th^ (>2 weeks later), at which point winds shifted back from the west—its prevailing direction—causing upwellings along the northern shore (Fig. S7D-E).

### Upwelling enriches for rare taxa by reshaping surface microbial composition

Spatial patterns of cell abundance varied across the lake surface in both May and September and correlated to surface temperatures (Fig. 1C). In May, cold, upwelling areas along the northern shore had lower cell counts while warmer offshore areas and the southern shore downwelling and Niagara-river impacted regions exhibited higher cell counts. In September, the upwelling zone at station 38 had the lowest cells abundances, whereas station 17, influenced by Welland Canal water, had the highest cell abundances.

Surface communities differentiated spatially, matching patterns in surface temperature cause by upwelling (Fig. 1D). Using UPGMA clustering based on the *GU^A^_0.5_* community dissimilarity, we defined 4 clusters in May (Fig. 1E) and September (Fig. 1F). In May, these clusters corresponded to the northern upwelling, an offshore pelagic group, the southern downwelling, and an isolated station near the Welland canal (Fig. 1D/E). Dispersion was also higher in surface May communities (Fig. S3C, PERMDISP, p = 0.002). In contrast, surface communities across September were comparatively homogeneous, with the Welland Canal also forming its own cluster alongside the station closest to Toronto. In contrast to May, however, September had outliers in the southern shore stations experiencing (or recently impacted) by upwelling (35 and 38, Fig. 1D/F, Fig. S3C). This differentiation was driven by two forces: (1) an influx of hypolimnion–associated taxa to the surface (Fig. S9A), likely due to mass effects, and (2) the appearance of numerous rare taxa not observed elsewhere in the lake during either season (Fig. S9B, S10A-D). Rare taxa were also comparatively frequent in May near the Welland Canal (Fig. S10B) but not along the northern shore upwelling (Fig. S10B).

### Environmental conditions shaped by stratification, not geographic distance, determine community composition

To address what ecological processes structured the spatial patterns shown in Fig. 1, we explored the relative importance of distance- and depth-decay in Lake Ontario’s microbial communities. Community similarity (1 - weighted UniFrac dissimilarity) was not correlated with geographic distance (Spearman Ranked R = -0.03, p = 0.44 in May and R = -0.05, p = 0.19 in September) but exhibited a strong relationship with depth in both months, particularly when the thermocline intensified in September (Fig. 2A-B, Spearman Ranked R = -0.50 in May and -0.65 in September, p < 0.0001 in both). Variance partitioning revealed that depth, month, and other environmental factors together explained 73% of the variation in microbial community composition (*GU^A^_0.5_*; Fig. 2C), a signal echoed in the PCA of environmental gradients (Fig. S3B). Geographic location, even when combined with environmental variables, accounted for only 3% of the variance.

**Figure 2.**
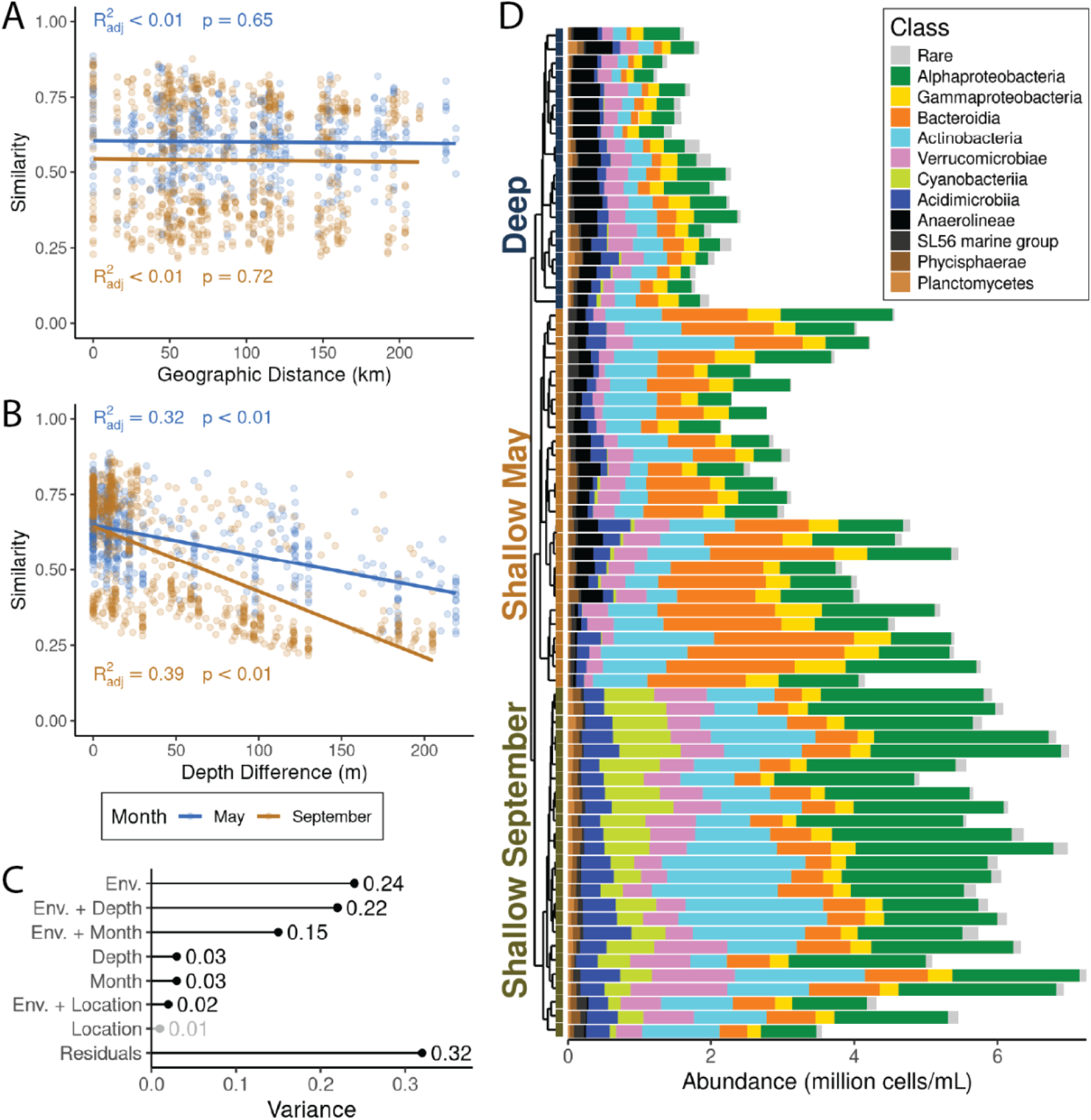
Thermal stratification—not geographic distance—drives microbial community structure in Lake Ontario. Microbial community (A) distance- and (B) depth-decay relationships by month using similarity defined as 1 - community dissimilarity (*GU^A^_0.5_*). (C) Variance partitioning of community dissimilarity. Environmental variables (Env.) corresponded to scaled physical and chemical data from each sample. Variables which were significant (all p < 0.001) when tested using an ANOVA-like permutation test for Constrained Correspondence Analysis post-dbRDA are in black whereas those that were not (i.e., location) are in light gray. (D) Community composition of all samples in absolute abundance, colored at the Class-level. Samples are arranged via hierarchical clustering as shown in Fig. S3A.

As a result of stratification, Lake Ontario bacterial communities were clustered into three distinct groups (Fig. S3A-C). These groups closely hew to sample depth (typically >30 m for deep samples, depending on thermocline depth) and month of collection, and are thus referred to as “depth-month groups.” This pattern was consistent across both hierarchical clustering (Fig. S3A) and principal coordinates analysis (Fig. S3C). Deep samples from both months grouped together (referred to as a single, “Deep” group), while shallow samples from May and September formed distinct clusters (p < 0.001, R^2^ = 0.55, PERMANOVA). Therefore, we categorized bacterial communities into three groups for analysis: Shallow May, Shallow September, and Deep (Fig. 2D). When comparing Deep samples between months, we include the month in parentheses, even though these samples clustered together (e.g. “Deep (May)”). Seasonal differences in sampling depth influenced the distribution of “mid” samples: in May, these were collected at the thermocline while in September they were taken at each station’s average depth, reflecting federal practices. Accordingly, “mid” samples from the thermocline were distributed throughout the Shallow May group (Fig. S3), whereas those from September were split between Deep and Shallow September groups depending on station depth and upwelling (Fig. S3).

Microbial cell abundances *across* these groups were strongly correlated with temperature (Fig. S11A; Spearman ranked R = 0.92, p < 0.0001). Abundances were highest in Shallow September, followed by Shallow May, and lowest in Deep samples (Fig. S11B). Within each group, cell abundances and temperature were correlated in Shallow May (Spearman ranked R = 0.771, p < 0.0001), but not in Deep samples (Spearman ranked R = 0.244, p = 0.299) or Shallow September (Spearman ranked R = 0.356, p = 0.0812). This may be because Shallow May also had the greatest temperate range (7.86 ℃ in Shallow May vs. 1.20 ℃ in Deep and 2.34 ℃ in Shallow September, excluding surface upwellings). Deep (September) samples also had fewer cells than Deep (May) samples (p = 0.031, Two-Sample Wilcoxon).

There was a core set of cosmopolitan taxa shared across all three groups, including the Alphaproteobacteria LD12, Actinobacteria acI lineages, Gammaproteobacteria LD28 and PnecB, and Verrucomicrobiae LD19 (Fig. 2D, Fig. S12B). Differentially abundant taxa included an enrichment of Chloroflexi in Deep samples, Bacteroidia in shallow May, and Cyanobacteria in shallow September (Fig. S12C)—including potentially harmful cyanobacterial taxa at 100,000 cells/ml, above the WHO limit for safe recreation (Fig. S13, [65]). A more thorough discussion of abundant microbial groups can be found in the Supporting Information.

### Season and depth modulate the strength of ecological selection across the lake

We deepened our analysis by quantifying community assembly processes within and between the three depth-month groups. Selection, dispersal, and drift were inferred by classifying the mechanism underlying species *turnover* between samples (Fig. S14). Although several frameworks exist for this purpose, they share a common logic: phylogenetic similarity between samples reflects the influence of selection, while taxonomic while taxonomic similarity reflects the influence of dispersal [56, 66]. Turnover not explained by either process is attributed to ecological drift. In this way, the relative importance of various assembly processes within one of between two groups can be estimated.

We applied iCAMP to quantify the relative contributions of five assembly processes: homogenizing and heterogeneous selection, dispersal limitation and homogenizing dispersal, and ecological drift. iCAMP integrates the phylogenetic similarity metrics β-Mean Pairwise Distance (βMPD) and β-Nearest Relative Index (βNRI) and taxonomic similarity metric Raup-Crick_Bray_ (RC_Bray_) [55]. Full details of the parameters and operation of iCAMP are available in the Supplemental Methods and Fig. S14, and discussed at length in [56].

Community assembly patterns differed across depths and seasons, with drift dominating deep samples, dispersal limitation increasing during stratification, and homogenizing selection prevailing in surface water. Drift dominated within Deep samples (58% in May and 44% in September), with homogenizing selection also playing a large role (34% - 38%, Fig. 3A). Drift accounted for almost half of pairwise turnover between deep May and September groups (49.2%) and between Deep and Shallow May samples (47%) but in September drift accounted for less variation between Shallow and Deep groups (16%; Fig. 3B). Dispersal limitation was also important in Deep (September) samples (19%) though not in Deep (May) samples (2%, Fig. 3A) and more pronounced when comparing Deep to Shallow samples, especially in September during stronger stratification (29% in September vs. 4.6% in May). Within Shallow samples, homogenizing selection was greater, especially in September compared to May (69% vs. 59%) (Fig. 3A). Across all depth-month groups, homogenizing selection explained 34%–69% of turnover, indicating consistent selection for a core microbial community throughout the lake (Fig. 3A).

**Figure 3.**
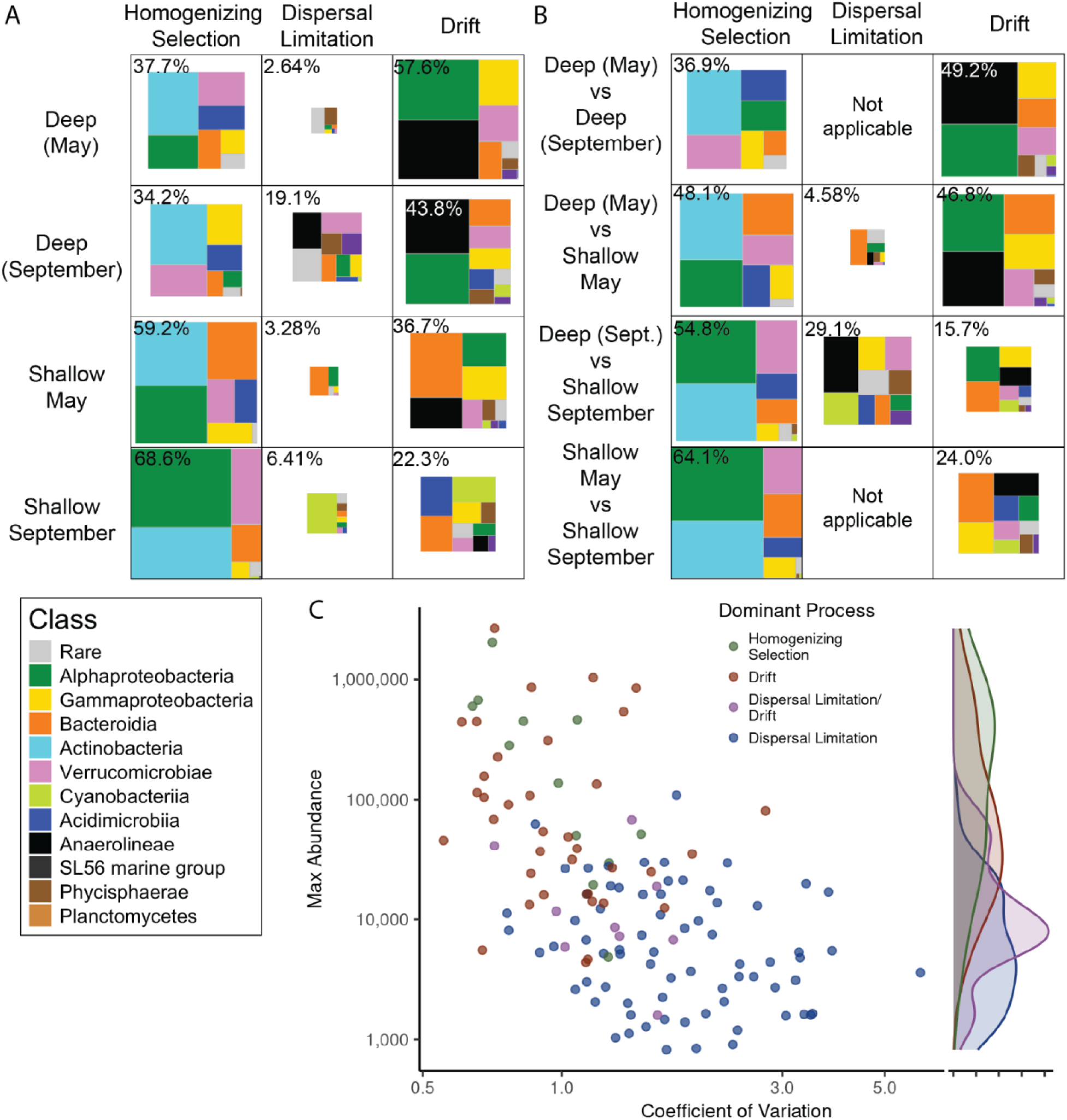
Contrasting forces of selection and drift structure microbial communities across depth. (A-B) Relative contribution of different taxonomic groups to microbial community assembly processes, as determined by iCAMP. The area of each box represents the relative contribution of that process (columns) to turnovers between samples (A) *within* a given depth-month group, or (B) *between* two depth-month groups. The percentage contribution of a given process to community turnover is in the top left corner, and boxes are scaled relative to the most influential process (*i.e.* homogenizing selection in shallow September at 68.6%), and the area represented by each Class is scaled relative to that Class’s contribution to that process. (C) The relative importance of community assembly processes within each phylogenetic bin, calculated by iCAMP. Absolute ASV counts were summed per bin per sample while the coefficient of variation was calculated as the standard deviation of abundances across samples by the mean abundance of that bin across all samples. The density plot on the right shows the distribution of each process across the maximum phylogenetic bin abundances. Three bins dominated by heterogeneous selection or a combination of drift and homogenizing selection were excluded.

iCAMP has an added benefit of quantifying assembly processes within smaller, monophyletic “bins” of closely related ASVs across the phylogenetic tree of a community (see Fig. S14). This improves accuracy of assembly quantification [56], and allows one to quantify the contribution of given taxonomic clades to assembly processes within each depth-month group (Fig. 3A-B), while also assessing the importance of assembly processes within a given clade (Fig. 3C). Homogenizing selection was primarily associated with the Actinobacteria, while drift was dominated by Alphaproteobacteria and Anaerolineae. Dispersal limitation between deep and shallow groups was primarily influenced by Bacteroidia in May and Anaerolineae and Cyanobacteria in September.

The contribution of taxonomic groups to community assembly was closely tied to their abundance within samples, particularly for within-group comparisons (compare Fig. 2C and Fig. 3A). More abundant bins had lower abundance variation across samples (Spearman ranked R = - 0.61, p < 0.0001) and were primarily shaped by drift and homogenizing selection (Fig. 3C). In contrast, low-abundance bins with high variance were dominated by dispersal limitation, while intermediate-abundance bins were influenced by both drift and dispersal limitation (Fig. 3C).

### Upwellings are an opportunity for novel microbial metabolic potential

Our analyses revealed that stratification structures microbial communities by modulating the relative influence of selection and drift. But what happens to microbial biogeochemical cycling when upwelling disrupts this physical structure? We used FAPROTAX to infer the abundance of genes associated with microbial metabolic functions (Fig. 4). FAPROTAX uses a manually curated database to assign metabolic functions to ASVs using their taxonomic assignment [57]. In comparison to other functional inference tools (like PICRUSt2), FAPROTAX only predicts metabolic functions which are conserved within a given taxonomic group, representing a more conservative prediction of the presence of a given function [67]. Upwelling zones exhibited a functional signature that blended traits from both the epilimnion (e.g., photoautotrophy; Fig. 4A) and the hypolimnion (e.g., ammonia oxidation; Fig. 4B). However, consistent with the presence of novel microbial taxa in the September upwelling zone (Fig. S9), we also detected functions that were unique to upwelling, including sulfate respiration (Figs. 1D and 4C) and methanotrophy (Fig. 4D). Sulfate respiration was assigned to ASVs in *Desulfuromonas sp., Sulfospirillum sp., and Sulfospirillum cavolei.* Methanotrophy was assigned to numerous genera within the Methylomonadaceae and Methylomirabilaceae. These results suggest that upwelling does not just remix existing microbial functional potential but rather creates entirely distinct microbial niches that may alter elemental cycling in the Great Lakes.

**Figure 4.**
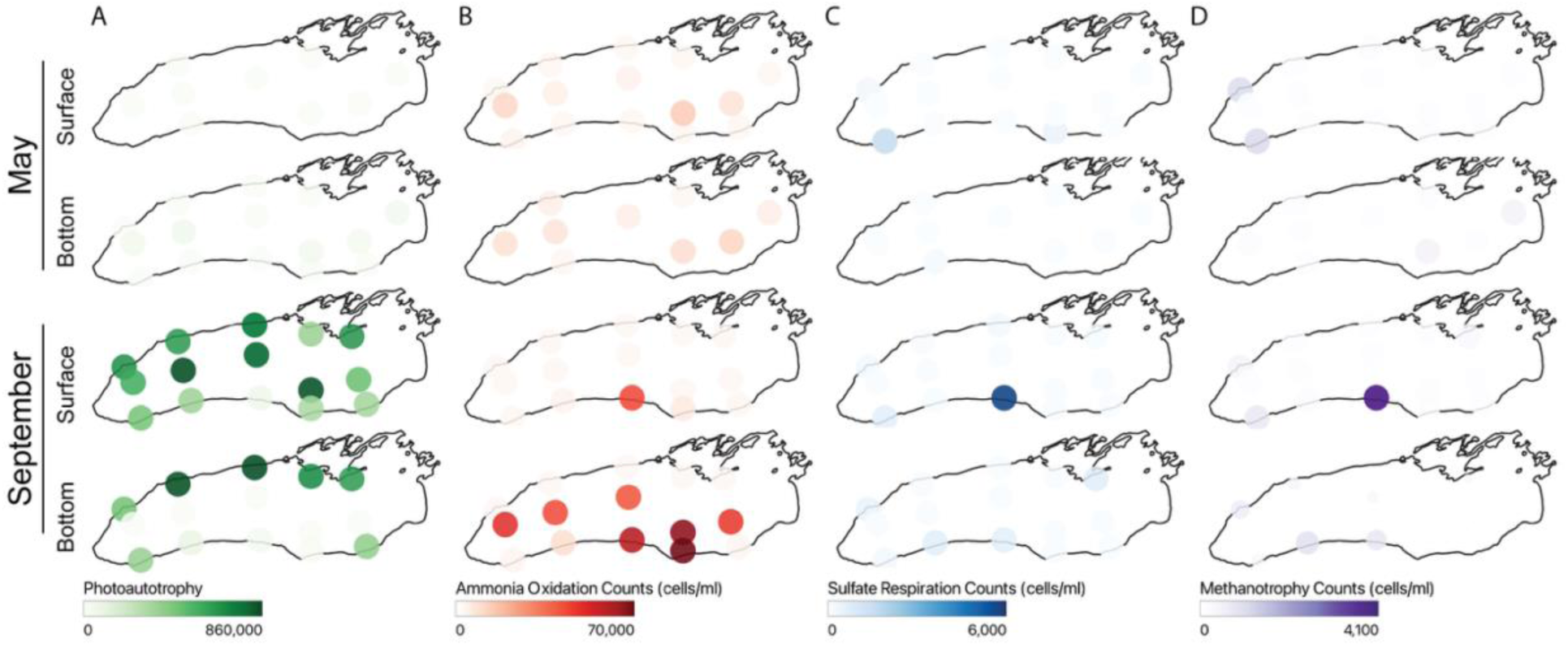
Upwelling creates distinct microbial functional potential. Functional traits were inferred using FAPROTAX [57] using the absolute abundance of each ASV as input, and results are shown for (A) photoautotrophy, (B) aerobic ammonia oxidation, (C) sulfate respiration and (D) methanotrophy. The color of each point is scaled by the inferred abundance of each functional trait, in units of predicted cells/ml. Note the range of values varies between functional traits.

## Discussion

### Physical forcing and ecological processes interact to govern microbial community structure

Microbial community assembly in Lake Ontario is fundamentally shaped by physical forces that structure the water column. Stratification imposed vertical gradients, with depth and season modulating the balance between deterministic and stochastic processes. Yet, when stratification was disrupted through upwelling, downwelling, or tributary mixing, community composition was rapidly restructured. These transient disturbances reveal how responsive microbial communities are to ephemeral but ecologically significant niches in large lakes.

### Upwelling creates transient but consequential microbial niches

Lake Ontario’s ocean-like circulation drives microbial turnover at spatial and temporal scales rarely quantified in freshwater systems. In spring, thermal bars trap nutrients and sculpt sharp near-shore gradients [68]. In summer, typical westerly wind events force Ekman transport, tilt the thermocline, and inject cold, nutrient-rich hypolimnetic water into the surface [27, 29, 32, 69, 70]. As winds relax, energy is released as internal waves: near-inertial oscillations offshore and coastally trapped Kelvin waves [26, 28]. Propagating anticlockwise at ∼60 km day⁻¹ [28], the Kelvin wave–jet complex drags the upwelling signal laterally and, as we show, sustains an easterly moving microbial disturbance that persisted over two weeks (Fig. S7D–E), a timescale relevant for microbial growth, succession, and metabolism.

Although physical limnologists have characterized these waves for decades, our data provide the first lakewide evidence that they restructure microbial communities via the mobilization of rare taxa. As winters have warmed, stratification in Lake Ontario now begins significantly earlier (1-3 months earlier) and lasts with greater intensity, extending the upwelling season and intensifying its ecological impacts [71]. Given that upwelling-generating winds occur every ∼10-12 days in Lake Ontario [30] and can persist for weeks, some segment of the shoreline is likely experiencing upwelling—and its associated microbial disturbance—at any given point during the stratified season. Upwelling-favorable winds have already increased by ∼45% over the past three decades [34] and are projected to intensify further [6], underscoring the urgency of linking physical forcing to microbial ecological response.

Despite their frequency, the microbial consequences of upwelling remain under-characterized, especially in freshwater systems [72, 73]. During the September event we tracked hypolimnetic lineages (e.g., Anaerolineae, Nitrospiria) surfacing at station 38 (Fig. S9A), which we estimate had coupled microbial and thermal signatures persisting for more than two weeks (Fig. S7D–E). Upwelling sites were uniquely enriched in microbial taxa associated with sulfate-respiration and methanotrophy (Fig. 4C–D), revealing that upwelling does more than mix water and microbes from the epi- and hypolimnion: it creates novel biogeochemical niches capable of rewiring microbial ecosystem function. As wind regimes intensify under climate change, resolving these physical–microbial linkages will be essential for predicting the function and resilience of Earth’s largest lakes.

### Rare taxa contribute disproportionately to community novelty in upwelling zones

Upwelling and tributary-influenced zones hosted microbial communities distinct from surface or bottom waters and were enriched in rare taxa (Fig. S9). In September, low-abundance hypolimnetic microbes dominated at the southern upwelling site. Richness was also highest in deep and shallow September (Hill ^0^D = 475 ± 93.5, Hill ^0^D = 473 ± 60.7, respectively, upwelling samples removed, Fig. S10D). However, when considering evenness, shallow September had the lowest microbial diversity (Hill ^2^D = 6.84 ± 1.89, vs. 10.2-18,2 in other groups). The stratified surface is shaped by strong homogenizing selection that limits diversity, but during upwelling this constraint is relaxed, allowing rare microbes to briefly establish and persist at low (yet detectable) abundances. In May, similar enrichment occurred near the Welland Canal but not in the upwelling zone—possibly due to colder temperatures at that station and consequent slower microbial growth rates [74].

Although our dataset captures spatial rather than temporal heterogeneity, the enrichment of rare taxa in upwelling zones may parallel the behavior of conditionally rare taxa. These are microbes that persist at low abundance until favorable conditions permit local proliferation [75]. Upwelling may therefore generate transient spatial opportunities analogous to the temporal pulses that activate conditionally rare taxa. These events allow rare lineages to emerge and contribute disproportionately to ecosystem function.

These rare taxa are often overlooked, but may serve as a latent reservoir of ecological innovation—a microbial seed bank whose functional potential emerges under transient, physically driven conditions [76]. The distribution of these rare taxa is most likely to be structured by dispersal limitation, so their presence may be both spatially and environmentally constrained. While the long-term consequences remain unquantified, our findings point to a cryptic but consequential microbial response. Upwellings act as drivers of microbial dynamism that move alongshore, persist for up to two weeks, and locally reshape ecosystem function in the world’s largest lakes.

Tributary inputs did not generate persistent nearshore–offshore gradients, in contrast to patterns in Lake Michigan [77]. Instead, strong offshore ecological selection filtered out rare taxa, reinforcing a cosmopolitan microbial core structured by drift and selection (Fig. 3). However, episodic enrichment near tributary inputs combined with physical forces including upwelling, downwelling and spring thermal bar formation (Fig. 1A, Fig. 4) may temporarily boost microbial diversity and function in nearshore zones. These findings suggest that rare taxa, though ephemeral, may play outsized roles in microbial ecosystem response, revealing an under-appreciated layer of adaptability in Great Lakes microbiomes.

### Depth and season drive a shift from selection to drift and dispersal limitation

Environmental gradients from stratification and hydrodynamics—not geographic distance— best explain microbial community structure, consistent with metacommunity theory and findings that dispersal is rarely limiting in aquatic systems [4]. Surface communities, especially in late summer, were shaped by homogenizing selection, yielding consistent assemblages across sites. In contrast, hypolimnion communities were more stable and governed by drift in both months and dispersal limitation only in September. The lack of dispersal limitation in the May hypolimnion suggests stronger hypolimnetic mixing (both laterally and vertically) than under strong stratification in September, though the seasonal strength of hypolimnetic currents in the Great Lakes is unclear [78, 79]. In May, depth-decay was weaker (Fig. 2B), and surface communities were more spatially variable (Fig. S3C) and similar to those at depth, likely reflecting ongoing niche differentiation after spring mixing. By September, stratification and low turbulence reinforced surface selection. These dynamics mirror trends in the Pacific Ocean, where surface microbes are selected and dispersal limitation intensifies with depth [80]. Similar patterns occur in oligotrophic lakes, where nutrient scarcity favors efficient, streamlined taxa [16, 18, 81], while productive systems show more stochasticity [82–84].

In the hypolimnion, stronger stratification constrained dispersal in September (Figs. 2B & 3A– B), and limited mixing fostered site-specific communities that were consistent across space and time (Fig. S3C), despite environmental stability. Drift was also the major community assembly force structuring hypolimnion communities, contributing 44-58% (Fig. 3A). Cold, oxygenated deep waters harbor slow-growing microbes, with low cell densities (Fig. S11B) [85], limited gene flow, and dormancy, all factors which weaken the strength of selection [86].

While cold, oligotrophic hypolimnia are often thought to be selection-dominated (*see Supplemental Discussion: Selection in the Hypolimion*; [24, 87–89]), our results align with recent ocean studies showing that drift increases with depth due to sparse biomass and isolation [80]. These effects were evident in Lake Ontario’s hypolimnion, where distance-decay was weak (Fig. S15E–F) but dispersal limitation was consistent. Rather than forming a continuous biogeographic gradient [20], deep microbial communities appeared as isolated parcels shaped by intermittent mixing, representing a spatial mosaic of stochastic processes.

### Microbial abundance and taxonomy are linked to assembly processes

Abundant taxa were governed by selection and drift (Fig. 3C), consistent with the idea that dominant microbes experience stronger ecological filtering [16, 22]. Rare taxa were more dispersal limited, reflecting restricted distributions, though it is unclear if this is a result of narrow niche breadth or a transient introduction from an external source [90]. Synthesizing multi-year datasets could help discern which rare taxa are transiently rare versus permanently rare.

Taxonomic identity also influenced assembly. Actinobacteria were primarily associated with homogenizing selection. Freshwater pelagic Actinobacteria (such as those in the *acI-acIV* lineages) are characterized by diverse metabolic capabilities and small genomes [91, 92]. In contrast, Alphaproteobacteria, dominated by the ubiquitous LD12 clade, were governed by drift. This contrast may reflect differences in ecological versatility. Widespread Actinobacteria may exploit more diverse niches via adaptive radiation [91]. In contrast, the distribution of Alphaproteobacteria like LD12 were structured by both selection and drift (Fig. 3). This may reflect stochastic variation in the relative abundances of these ubiquitous taxa; while their ubiquity reflects a history of selection or dispersal advantages, variations in their distribution in Lake Ontario at a given timepoint appears stochastic. This highlights the importance of drift in structuring microbial distributions, especially for highly abundant taxa.

These patterns underscore that community assembly cannot be explained by abundance alone. Phylogenetic identity, ecological function, and life history traits all shape how taxa respond to environmental gradients and physical forcing. Yet, our interpretation is limited by the temporal resolution of our sampling, which captured only two stratified timepoints. Key transitions, such as thermocline formation, fall mixing, and inverse stratification-remain unresolved [7, 93]. Finer habitat delineations unexplored here (such as shorelines, benthic surfaces, and suspended particles) are also likely to influence assembly processes [77, 94, 95]. Finally, while iCAMP offers valuable insights, its inferences reflect model assumptions and should be applied with appropriate caution (*see Supplemental Discussion: Biases in iCAMP*).

### Upwelling-driven microbial restructuring alters functional potential

Hydrodynamic restructuring of microbial communities carries clear biogeochemical consequences. Upwelling, tributary mixing, and downwelling redistribute microbial taxa and their metabolic potential, creating biogeochemical hot spots that are both spatially and temporally dynamic [32, 32]. The chemical and physical environment of the upwelling generally represents a mix of deep and shallow water (Fig. S16). There is modest enrichment in total nitrogen and NH_4_. We did not observe a difference in sulfate concentrations, despite an enrichment in predicted sulfur respiration, which could be due to cryptic cycling (Fig. 4, Fig. S16) [96]. Our results, however, suggest that upwelling introduces rare, functionally distinct taxa into the surface waters, including lineages associated with sulfur cycling, methane oxidation, and ammonia oxidation, potentially shifting biogeochemical processing. These lineages may be at abundances below detection throughout the lake, but the unique combination of light, warmth, and chemical composition within the upwelling provides an opportunity for their growth. Testing this hypothesis would require sampling an upwelling with greater temporal resolution, in addition to bottle experiments where upwelling waters are exposed to different light and temperature regimes. These changes likely unfolded over weeks, as Kelvin wave–driven upwelling swept laterally along the nearshore, suggesting that microbial metabolism responds rapidly to these common physical forcings.

Functional inference based on metabarcoding, however, has inherent limitations. The 16S rRNA V4 region lacks the resolution to capture fine-scale genomic or functional differences, and strains or species with identical 16S V4 sequences (i.e. identical ASVs) may differ in metabolic capacity [97, 98]. Additionally, variable gene copy numbers and high dormancy rates in freshwater microbes can decouple gene presence from metabolic activity [67, 99]. As a result, the accuracy of functional inference depends on the trait of interest, the environment, and the size and curation of reference databases [57, 98, 100, 101]. For instance, nitrification rates in Lake Erie have been shown to be uncoupled from nitrifier abundance [102, 103].

To stay conservative, we used FAPROTAX, a relatively stringent approach; only 24% of ASVs matched at least one annotated function, likely underestimating the true functional potential of these communities. Future work using metatranscriptomics could more directly quantify metabolic responses to upwelling [99, 104, 105]. Ideally, these inferences would be validated by experimental incubations that measure microbial biogeochemical rates in upwelled waters [106–108].

### Microbial biogeography emerges from physical-ecological coupling

Microbial communities in Lake Ontario are not static, but dynamically assembled by the interaction of physical forcing and ecological processes. As northern lakes warm, deepening and lengthening stratification will isolate hypolimnetic communities [109–111] while enhancing surface selection, reshaping microbial biogeochemical cycling in ways we are only beginning to quantify. At the same time, the stratified (and therefore upwelling) season is lengthening with climate change, while upwelling-favorable winds are becoming more frequent and are predicted to intensify, increasing the prevalence and impact of microbial disturbance zones alongshore [6, 34, 71].

Our data show that such disturbances are not rare—recurring 2-3 times a month-with their influence persisting for weeks. These physical pulses are biologically important as they override stratification and assemble novel shoreline microbial communities, remodeling ecosystems on meaningful ecological timescales.

By pairing high-resolution microbial, ecological, and physical limnology data, we demonstrate that community structure and function in large lakes are governed not just by who arrives or survives, but also by how water moves. These systems offer natural laboratories for testing metacommunity theory, bridging freshwater and marine paradigms, and forecasting how microbial life will shape and respond to a changing climate.

## Acknowledgments

We thank Eric Osantowski, Matthew Pawlowski, Anne Scofield, Ben Alsip, and the science staff in the Great Lakes National Program Office of the US EPA and the captain and crew of the *R/V Lake Guardian* for facilitating sample collection. We thank Evie Brahmstedt, Sophia Aredas, and Sophia Richter for support in September field work. We thank Linda Cote and the BRC Genomics Facility (RRID:SCR_021727) at the Cornell Institute of Biotechnology for sequencing experiments. We are grateful to Andrea Giometto and his lab for flow cytometry support. We appreciate the feedback from Joe Atkinson and also the EPA & NOAA Great Lakes Modeling Group led by James Pauer. Thank you to the anonymous reviewers for their valuable feedback.

