## Supporting_Information for "Upwelling periodically disturbs the ecological assembly of microbial communities in Lake Ontario"

### Supplemental Methods

*Sample collection*

Depth profiles were collected as the rosette/CTD descended the water column and water samples were collected as the CTD ascended. Samples were collected without regard to time of day (though Stn 33 was sampled both night and day in September). As such, bottom samples were always filled first, and immediately taken to the lab for filtering. For all samples, 5 mL of water was initially sub-sampled for flow cytometry samples and kept in the fridge. First replicates at each depth were filtered immediately, while second replicates were kept refrigerated until the first replicate was done. Water samples were taken in duplicate from separate niskins and pre-filtered consecutively through sterilized 200 $\mu$m and 20 $\mu$m Nitex mesh (Wildco) into sterile 10L bottles (Nalgene). Bottles were rinsed twice with sample water before filling. Microbial samples were filtered using MasterFlex Peristaltic Pumps onto a polyethersulfone 0.22 $\mu$m filter (MilliporeSigma). Samples were filtered for a maximum of 25 minutes, with flow through volumes ranging from 1500 mL to 8760 mL. Filters were flash frozen in liquid $N_2$ and stored at -80C.

*Environmental data*

Water chemistry and chlorophyll a data were generated by the U.S. Environmental Protection Agency's Great Lakes National Program Office according to their standard protocols [1]. Data included $NH_4$, $NO_x$, Soluble Reactive Phosphorus, Total Nitrogen, Total Phosphorus, K, Na, Ca, Mg, Cl, $SO_4$, Dissolved Organic Carbon, Si, and Chlorophyll-a. Data was downloaded from the Great Lakes Environmental Database system, which can be accessed through the Environmental Protection Agency Central Data Exchange (https://cdx.epa.gov/). Temperature profiles from the Great Lakes Acoustic Telemetry Observation System (GLATOS) moorings were collected and provided by the Ontario Ministry of Natural Resources, Fisheries and Oceans Canada, the U.S. Fish and Wildlife Service, U.S. Geological Survey, and Queen's University, supported in part by the Great Lakes Observing System (GLOS). Data are publicly available through the GLOS Seagull platform (https://seagull.glos.org/data-console/groups/379). Lake Erie water temperatures at the Niagara River in Buffalo were downloaded from the National Weather Service (NOAA, https://www.weather.gov/buf/LakeErieMay)

*Microbial Sampling*

Duplicate microbial samples from separate niskins but the same CTD/rosette cast were immediately processed on board after water collection. For the first replicate, there was a maximum time of 96 minutes and a median time of 35.5 minutes from the Niskin bottle being fired to the sample being frozen. Between replicates of the same station and depth, 1 L of MilliQ was used to rinse the filtration tubing. Between stations, sample bottles and filtration tubing were sterilzed by rinsing and incubating at room temperature for 10 minutes with 10% (v/v) sodium hypochlorite and then rinsed three times consecutively with MilliQ 18.2 MΩ water. A negative control was prepared on the last day of each cruise using the same tubing sterilization procedure and running 4 L of MilliQ 18.2 MΩ water through a 0.22 $\mu$m filter. Read depth or richness was not correlated to time from sampling till freezing, water extracted, or DNA concentrations.

*DNA extraction & Illumina sequencing*

All DNA extractions were carried out using the Qiagen DNeasy PowerWater kit. Briefly, samples were incubated at 65C in PW1 for 5 minutes, vortexed for 5 minutes, centrifuged at 4,000g for 1 minute, again incubated for 5 minutes, and vortexed for 5 minutes. Otherwise, extractions were carried out as per manufacturer directions, and eluted with 40-50$\mu$l Elution Buffer. DNA yields were quantified using the dsDNA Broad-Range kit from Qubit and ranged from 0.302 ng/µL to 151 ng/µL. 149 samples were sequenced on an Illumina MiSeq using v2 chemistry with 2x250 paired end reads, 10% underloading and 20% PhiX spike-in. Cluster density was 692, with 15.72 million raw reads and 13.55 million reads passing filter and an average of 105.5 thousand reads per sample.

*Microbial bioinformatic processing*

Raw Illumina sequences were processed into amplicon sequence variants (ASVs) using the standard workflow in dada2 package in R [2]. Primers were trimmed and sequences were quality-controlled using the dada2::filterAndTrim command with truncLen = c(240,220) and trimLeft = c(20,20) with maxEE = c(1,1) and maxN = 0, based on quality plots. Merged ASVs were length filtered to 252 or 253 bp. A median of 64.1% of reads were retained.

Taxonomy was assigned using both the TaxAss freshwater database and the Silva 16S rRNA databases [3, 4]. If ASVs were classified using the curated freshwater taxonomy, that classification was retained; otherwise, the Silva classification was used. For the freshwater taxonomy, ASVs were classified using the RunSteps_quickie.sh script using the FreshTrain15Jun2020silva138 database with default parameters [3]. For the Silva database, ASVs were classified using dada2::assignTaxonomy function, against the Silva non-redundant (99%) v138 database and dada2::addSpecies function using the Silva species v138.1 database [4].

*Measuring community assembly processes with iCAMP*

iCAMP quantitatively estimates the importance of various assembly mechanisms using measures of phylogenetic and taxonomic dissimilarity. As assembly proccesses likely vary within sub-populations of a given community, iCAMP first divides the community into smaller, monophyletic lineages, in which the relative importance of assembly processes are evaluated Fig S14). Using our scaled environmental parameters (Fig. S3B), we tested the average phylogenetic signal present in bins across bin.size.limits (24 and 48) and ds values (0.1, 0.3, and 0.5) using

Mantel Tests as performed with the ps.bin function. The average correlation between phylogenetic distance and environmental niche (MeanR) was highest with smaller bin sizes (Fig. S5). As ds played a small role in the MeanR value, we opted for a ds limit of 0.5, opting for slightly fewer, larger bins. Over-all findings (that homogenizing selection and drift dominated community assembly) remained consistent at across bin size limits and ds values (Fig S5).

Selective processes were quantified first using $\beta$ Net Relatedness Index ($\beta$NRI) and then dispersal processes were quantified using a modified Raup-Crick (RC) method based on the Bray-Curtis dissimilarity of relative abundances [5, 6]. However, after testing for the normality of null distributions (null.norm), most bins had non-normal null distributions, so a one-tail non-parametric confidence test was instead used, by using the change.sigindex function with argument Confidence. Significance of inferred assembly processes between each month/depth group was tested using the icamp.boot function with 1000 iterations; all comparisons shown in Fig. 3A were considered significant.

The relative importance of an assembly processes in explaining the turnover between two samples is then estimated as the sum of bins dominated by that process, weighted by those bins' abundance overall. To divide the relative contribution of each microbial Class to a given process, we accessed the Class.maxNamed column in the Bin.TopClass slot.

To estimate a bin's abundance and coefficient of variation, the absolute abundance of all ASVs in each bin was summed for each sample. The coefficient of variation was calculated by dividing the standard deviation of absolute abundances across samples by the mean absolute abundance of that bin across all samples. To facet bin-level processes between Phyla, bins were divided based on Top.TaxonPhylum, and labeled by Top.TaxonGenus if available; otherwise, the lowest taxonomic rank available was used.

### Supplemental Results and Discussion

*Microbial Community Composition across the thermocline*

Microbial communities in Lake Ontario shared a core set of cosmopolitan taxa, but certain Classes were closely tied to specific depth-month groups. Alphaproteobacteria (predominantly LD12, a close relative of the marine SAR11 lineage), Actinobacteria (including acI-B1, acI-C2, acI-A7, acI-B1, acI-A6, acSTL-A1, acI-A5, and acI-A3), Bacteroidia (such as unknown species and bacI-A1, bac-IIA, and bacIII-A), Gammaproteobacteria (LD28 and PnecB, formerly Betaproteobacteria), and Verrucomicrobiae (mostly LD19, SH3-11, and unknown species) were ubiquitous across all samples (Figs. 2D & S9A-B).

Differential abundance analysis revealed distinct habitat preferences among taxa. Chloroflexi Anaerolineae, including an ASV sharing 99.6% identity with CL500-11 from Crater Lake [7], were enriched in Deep and shallow May samples (Fig. S12). Deep samples had elevated levels of nitrifiers, including the ammonia-oxidizing Archaea Nitrososphaeria and the nitrite-oxidizing Bacteria Nitrospiria. In contrast, Bacteroidia were more prevalent in shallow waters, with the largest populations in May. Acidimicrobiia and Cyanobacteria (mostly *Cyanobium PCC-6307*) were prominent in shallow September, along with two potentially harmful cyanobacterial genera (Fig. S13A-B). *Dolichospermum NIES41* was almost 1% of the community (>60,000 cells/mL) at station 17 near the Welland Canal (Fig. S13C), while *Microcystis PCC-7914* was abundant across multiple stations, peaking at ~100,000 cells/mL at station 43 near Cobourg, Ontario (Fig. S13D). As these samples were pre-filtered through a 20$\mu$m mesh before sequencing, these values could underestimate the true abundance of these colonial cyanobacteria.

*The Upwelling Niche*

It is notable that Great Lakes upwelling events in a given location tend be ephemeral, lasting a few to several days and create concomitant downwelling events across the lake (Fig. S7, [8]). These upwelling events create pulses of nutrients and bottom-dwelling taxa which disturb the dispersal barrier created by the thermocline. However, upwelling events can propagate laterally as slow-moving Kelvin waves, and occur regularly (*e.g.* a few per month) in stratified Lake Ontario (Fig. S7, [8, 9]). In September, upwelling stations disproportionately featured unique taxa not observed elsewhere (Fig. S9). Accordingly, upwelling zones represent a consistent, if emphemeral, niche in Lake Ontario with unique microbial diversity,

*Selection in the Hypolimnion*

While selection was less influential in shaping the microbial hypolimion community, selection in deep waters is likely intense for microbes who actively grow in this environment, especially over long time scales. For example, biogeographic patterns of Chloroflexi C500-11, a ubiquitous and (presumably) active member of the hypolimnion, suggest strong purifying selection under hypolimnion conditions [10–12]. Similar results have suggested globally distributed freshwater lineages of nitrifying taxa, whose occurrence is almost exclusively hypolimnetic due to photoinhibition of nitrification [13]. Multi-annual time series or sequencing approaches which differentiate active from dormant taxa in the hypolimnion may make homogenizing selection more influential.

*Rarity in Lake Ontario*

While the distribution and assembly of rare bacteria are often more stochastic (see [14]), different assembly mechanisms can produce unique "types" of rarity [15, 16]. Four types of rarity have been proposed: permanently rare, permanently rare with periodic distributions, transiently rare, and conditionally rare [16]. Because we only sampled two time points, we cannot directly assess conditionally rare taxa, which are defined by temporal fluctuations in abundance. However, we did observe taxa with low overall prevalence but high maximum abundance (Fig. S12A). These patterns could reflect conditional rarity in a temporal sense, or a spatial analog in which taxa thrive only under isolated environmental conditions in one area of the lake.

Capturing microbial communities in Lake Ontario at high temporal and spatial resolution would be required to distinguish between temporal and spatial conditional rarity; this sampling design would also be difficult in practice. For the taxa that are rare in our dataset, drift and dispersal limitation emerged as the dominant assembly mechanisms, suggesting most are either either consistently rare with periodic distributions or transiently rare [16]. This pattern is consistent with the absence of a distance-decay relationship among rare taxa, as shown using the abundance-unweighted UniFrac dissimilarity (Fig. S15C and S15E). Together, these results indicate that dispersal limitation signals in rare bacterioplankton likely reflect drift-driven differentiation across temporarily separated parcels of water, rather than linear range dispersal limitation [17].

*An expanded picture of microbial diversity in Lake Ontario*

Previous studies of Lake Ontario microbial diversity focused on a few offshore stations [18–21] or highly impacted embayments [22]. Our findings confirm dominant taxa like the *acI* and *LD12* lineages throughout the lake, the oligotrophic heterotroph *CL500-11* in the hypolimnion, and non-toxic *Synechococcus* in summer surface waters. Notably, we observed an abundance of Bacteroidia (*Flavo-A*, *bacI*, and *bacII*) in spring and a previously unreported diversity of Acidomicrobiia (*acIV* lineages like *Iluma-A1*), expanding known microbial diversity of the lake. Though, these differences may reflect primer bias or the effects of a different pre-filtration strategy (*see Supplemental Discussion: Biases in Sampling and Sequencing*). Filamentous Bacteroidia may dominate in spring by averting grazing [23, 24]. Additionally, we detected the *acI-C2* lineage, characteristic of the lower Great Lakes [18], which was significantly more abundant in shallow September (Fig. S17), suggesting niche selection to the relatively warmer lower lakes [25].

*Biases in iCAMP*

These data, while novel in spatial breadth, rely on discerning patterns rather than directly measuring processes and depend on statistical assumptions, such as niche partitioning being phylogenetically conserved and dispersal being random [5, 6]. However, dispersal, though modeled as a stochastic process, can be determined selectively for a given microbial clade [26, 27]. As selection is evaluated first by iCAMP, this approach may bias results toward detecting selective processes, potentially underestimating dispersal's role in community assembly [6]. We have good reason to use iCAMP, which includes the phylogenetic binning step, over previous, simpler tools that consider the community as a whole (e.g. QPEN, [5]). As we increase the bin.size.limit, our analyses become more generalized across the entire community, "smoothing" potential variations in assembly processes, and effectively weighting our analysis of assembly processes towards increasingly abundant lineages. When we omit any phylogenetic binning (e.g. using metrics originally proposed in Stegen 2013, rather than iCAMP), homogenizing selection becomes almost exclusively dominant (Fig. S14B).

*Biases in Sampling and Sequencing*

Choices of pre-filtration and sequencing strategy are consequential to the measure of microbial diversity. Even at 20 µm pre-filtration, many particle-associated, colonial, and filamentous bacteria are likely removed from our dataset. In addition, all primers used in metabarcoding amplification will show some bias [28]. These two factors could explain some of the differences in our dataset compared to [18]. Their more stringent pre-filtration at 1.6 µm may have reduced the abundance of large or filamentous taxa, like the Bacteroidetes and Cyanobacteria (outside of *Cyanobium*). In addition, they tested both V4 and V4-V5 16S rRNA region primer sets, and opted to use the V4-V5 as it provided greater genomic resolution to differentiate taxa. As such, we may have underestimated the abundance of Alphaproteobacteria *LD12* and Actinobacteria *acI-B*, while the V4-V5 primer set seemed biased against Acidimicrobiia *acIV* lineages, which were prominent in our dataset.

In addition, while our dataset is a valuable tool to probe "bottom-up" controls on microbial dynamics, but we did not consider important "top-down" controls through zooplankton grazing, which may be particularly relevant in oligotrophic lakes [29, 30]. While simultaneously collected zooplankton samples were collected during the cruises, the methods used to collect them likely removed the smaller taxa who are the predominant microbial grazers like ciliates. DNA approaches like 18S rRNA metabarcoding on non-prefiltered samples would provide valuable insights into the relationships between micro-eukaryotic and prokaryotic plankton.

### Supplemental Figures

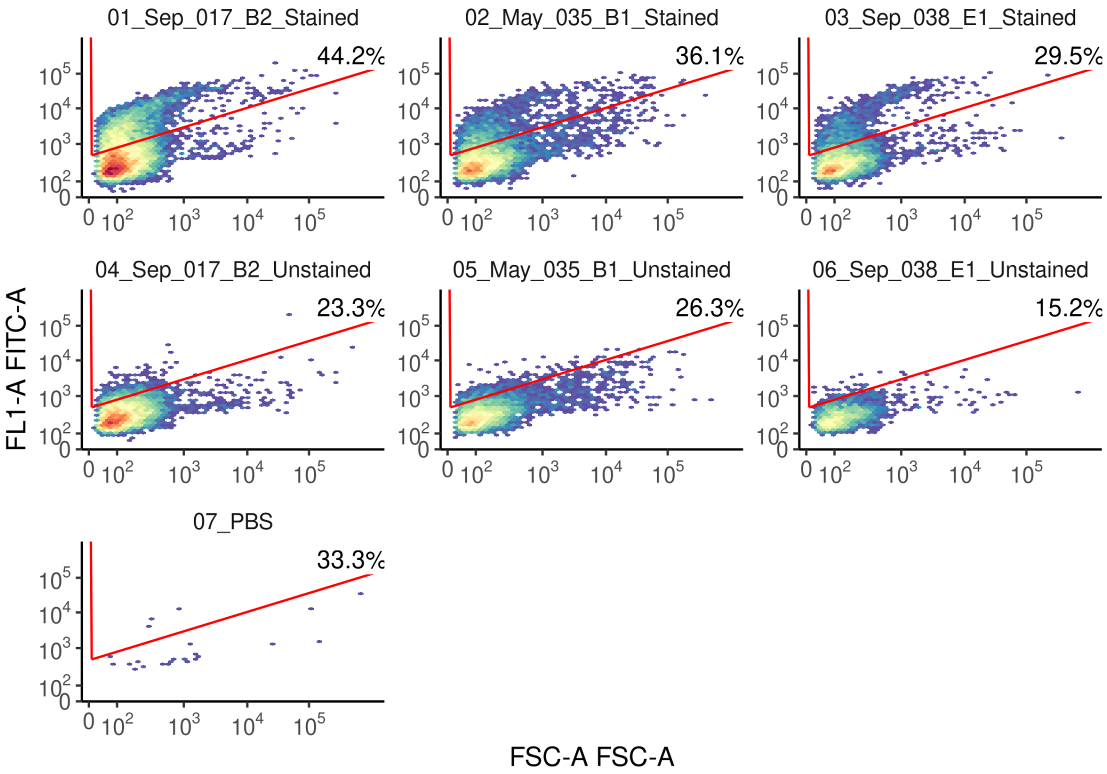


*Figure S1. Example Gating for SYBR-Positive Events.* Density plots generated showing green fluorescence (with the blue laser FL1 filter on y-axis) versus forward scatter (FSC on x-axis) for three samples stained with SYBR green (top row) versus unstained (middle row) and a PBS control (bottom row). Raw events were filtered with a FSC-H filter of 100 and FL1-H filter of 400. The polygon gate which defines positive events (cells) is available within the Github repository MarschmiLab/Pendleton_2025_Ontario_Publication_Repo/data/04_cytometry_exports/fcs_files/ .

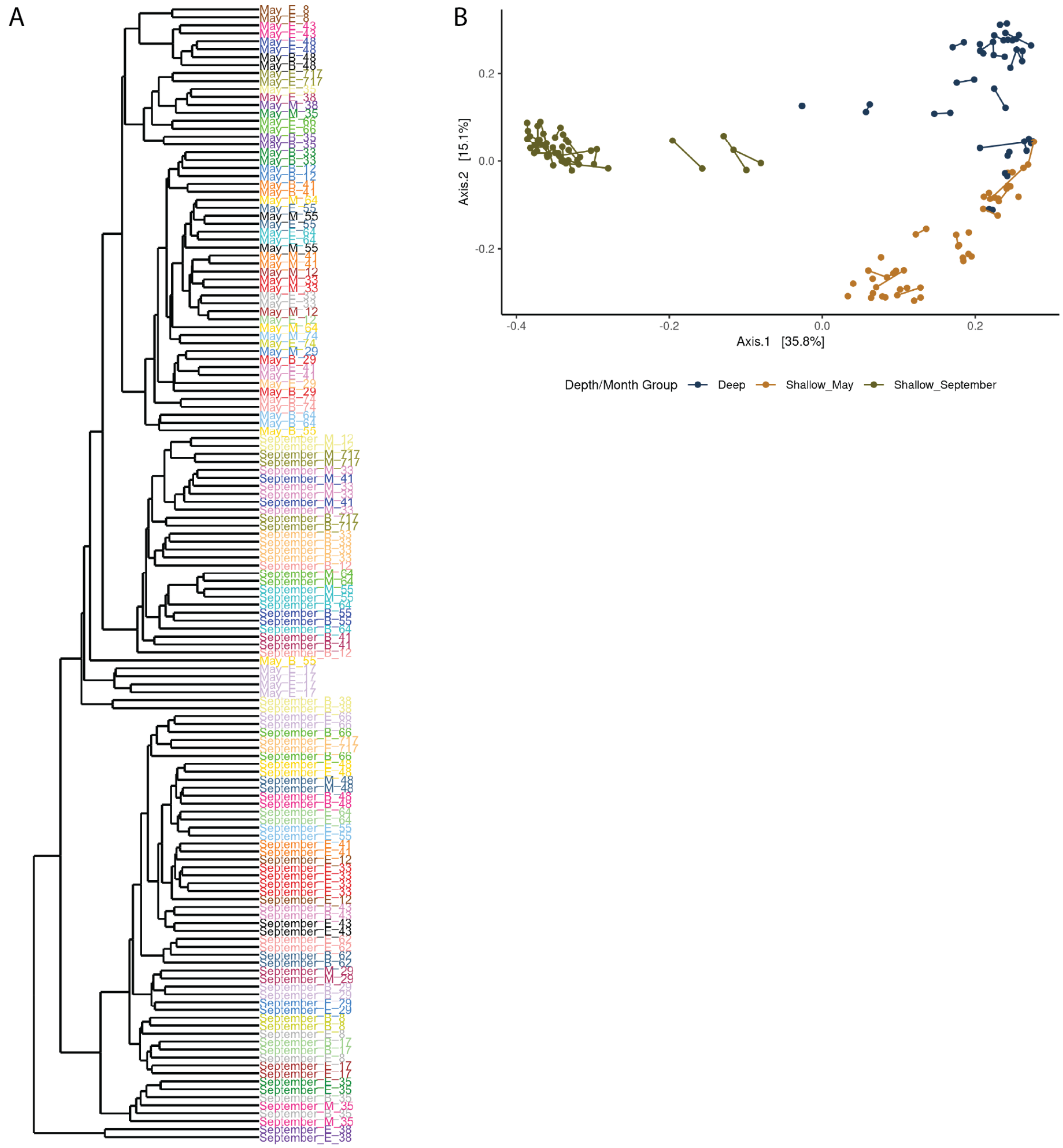


*Figure S2. Replicate samples had highly similar composition and were merged for following analyses.* (A) Hierarchical clustering of all replicate samples using the abundance-unweighted version of the Bray-Curtis dissimilarity, also known as the Sørensen Dissimilarity. Replicates of individual samples share colors; most replicates are highly similar to each other. Sample names correspond to Month_Depth_Station, where Depth is "E" for surface (epilimnion) samples all taken at 5 m, "M" for mid samples (thermocline or middle of water column), and "B" for bottom samples for 2m above the benthos and therefore varies based on total depth. These depths represent the standard U.S. and Canadian federal practices and is therefore consistent with historical samples from these stations. (B) Principal Coordinates Analysis of all replicate samples using the Sorensen distance. Replicates are colored by their depth-month groups and connected via a line. In assessing replicate similarity, we used Sørensen Dissimilarity as it is sensitive to changes even within rare taxa, even though an abundance weighted generalized unifrac metric is used throughout the rest of the manuscript (see methods, *Ecological Statistics*).

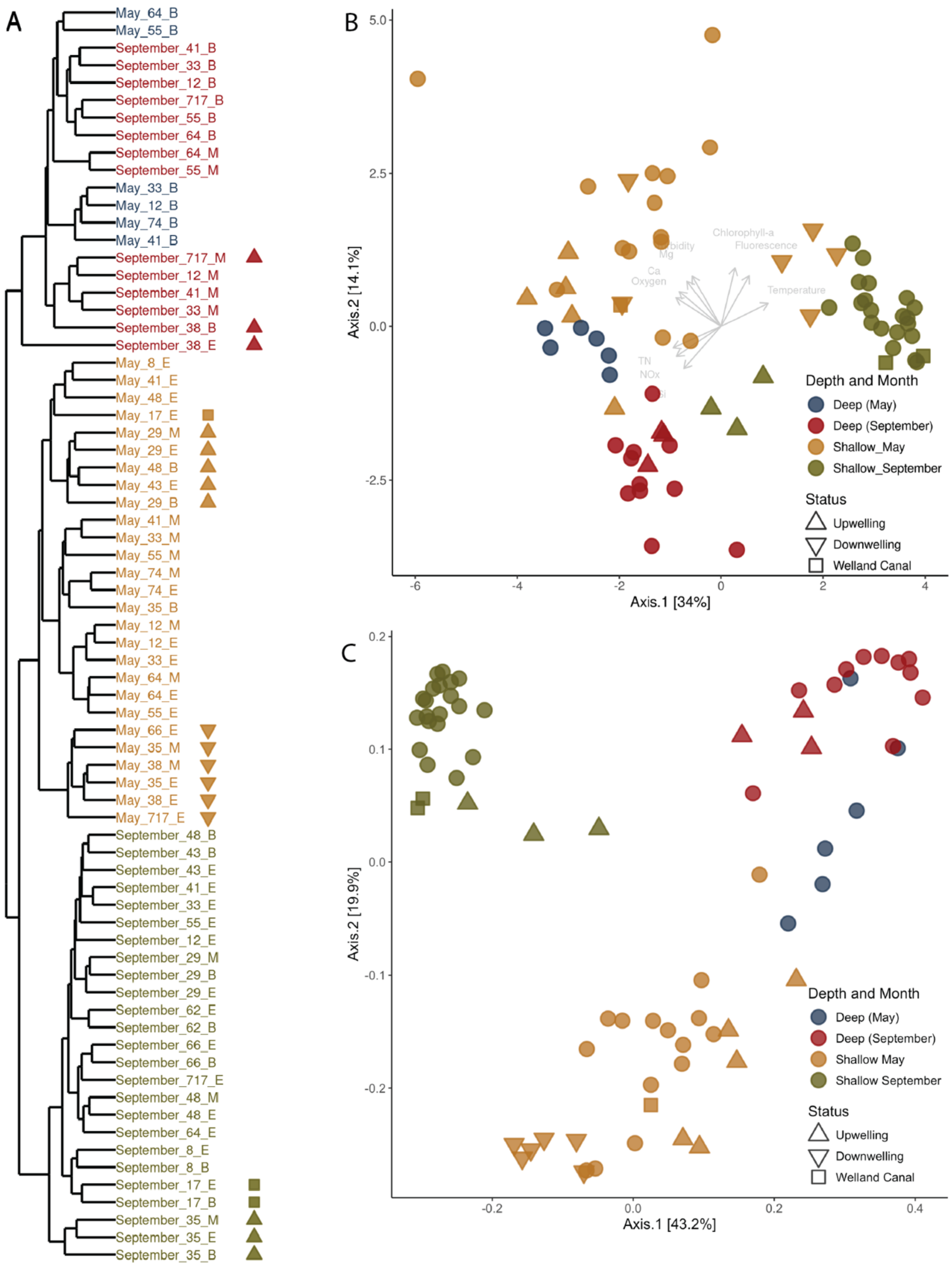


*Figure S3. Depth-Month Group Splits* (A) Month and depth groups were defined with the abundance-weighted UniFrac dissimilarity using a UPGMA hierarchical clustering method, cutting the tree at 3 groups, though for illustration deep samples from May and September are differentiated by color. Symbols denoting upwelling and downwelling stations and the Welland Canal are consistent across all three panels. Sample names correspond to Month_Depth_Station, where Depth is “E” for surface (epilimnion) samples, “M” for mid samples (thermocline or middle of water column), and “B” for bottom samples. (B) PCA of physical and chemical parameters collected via CTD sensors and analyzed by the EPA. All variables were scaled and centered before ordination. Only variables assessed as significant via vegan::envfit ($R^2>0.5$, $p < 0.05$) are included. (C) PCoA of microbial community data, using the abundance-weighted UniFrac distance as input, calculated with non-normalized absolute abundances of each ASV. Shallow May had significantly higher dispersion than shallow september (PERMDISP, $p = 0.002$).

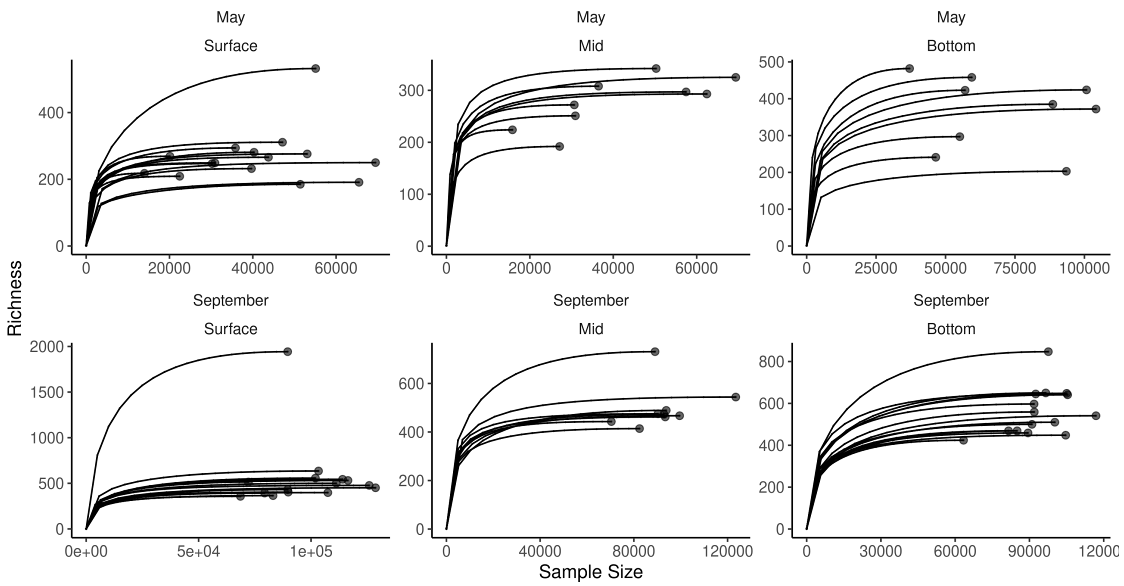


*Figure S4. Rarefaction curves of merged samples.* Rarefaction was performed using iNEXT on merged replicate samples sub-sampling at forty "knots" within each sample. Curves terminate at the observed richness.

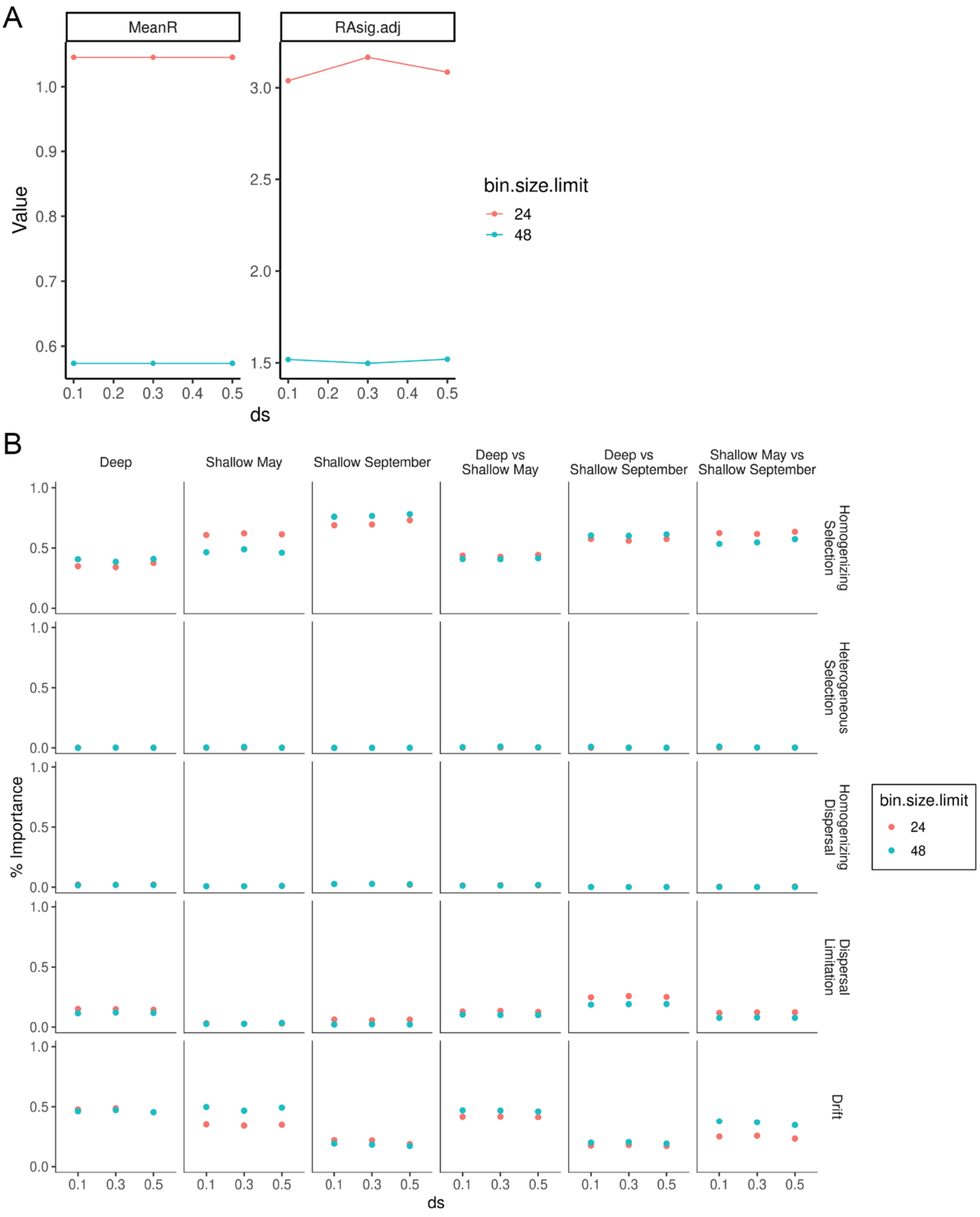


*Figure S5. iCAMP results across ranges of paramterers ds and bin.size.limit.* Two parameters control the phylogenetic "binning" in iCAMP; the initial phylogenetic distance to cut the tree (*ds*) and the minimum number of ASVs in each bin (*bin.size.limit,* Fig. S14). It is recommended to tune this parameters to maximum the phylogenetic signal of niche preference within bins (the correlation of ASVs' abundance to scaled environmental parameters). (A) Using our scaled environmental parameters (Supporting Information Fig. S3B), we tested the average phylogenetic signal present in bins across *bin.size.limit* (24 and 48) and *ds* values (0.1, 0.3, and 0.5) using Mantel Tests implemented in iCAMP::ps.bin function. MeanR represents the average correlation (R) between phylogenetic distance and environmental niche (R), and RAsig.adj corresponds to the number of comparisons where R was significantly different from zero. (B) Using a reduced number of iterations (100), we re-ran iCAMP across a range of *ds* and *bin.size.limit* values to compare the estimated importance of assembly processes both within and across depth-month groups.

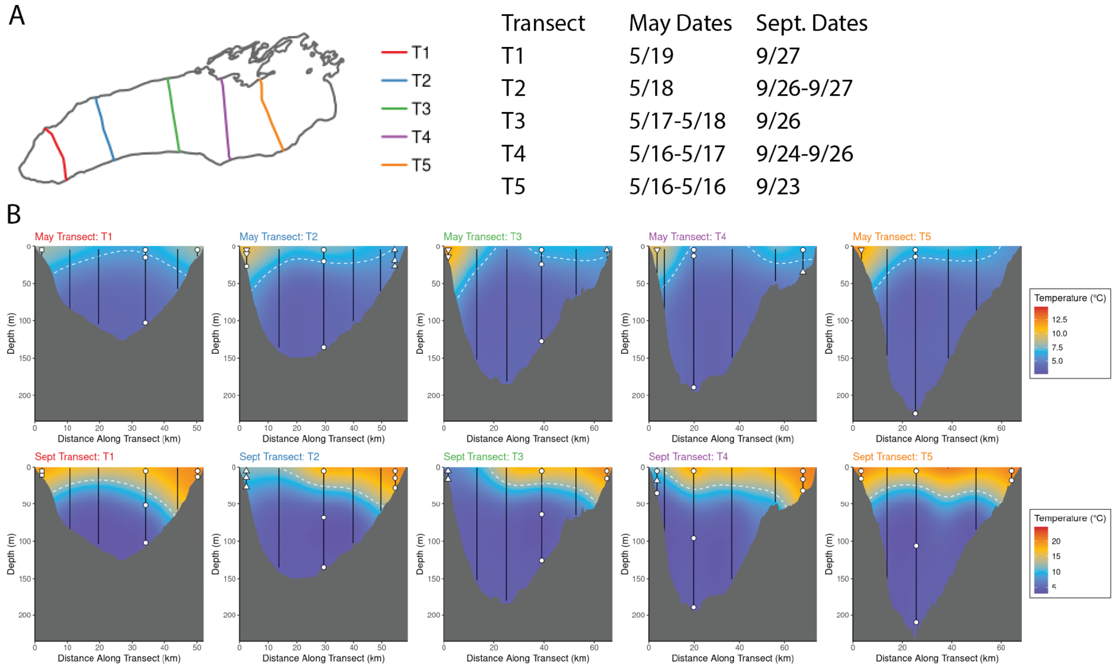


*Figure S6. Temperature Cross Sections of Lake Ontario in May and September* (A) Transects are labeled 1 through 5, moving eastward. (B) Temperature data from CTD casts was interpolated between stations along a transect using multilevel B-splines in R across a 300x300 step grid for each transect with 5 hierarchical levels. The grey bottom reflects bathymetry of the lake, moving along the transect from the southern shore to the northern shore. A 6°C isotherm is included in May, and a 12°C isotherm in September. Sampling locations are noted via white symbols; upwelling stations are upward-facing triangles, downwelling stations are downward-facing triangles, and the station nearest the Welland Canal is a square. Note that temperature scales are different between May (top row) and September (bottom row).

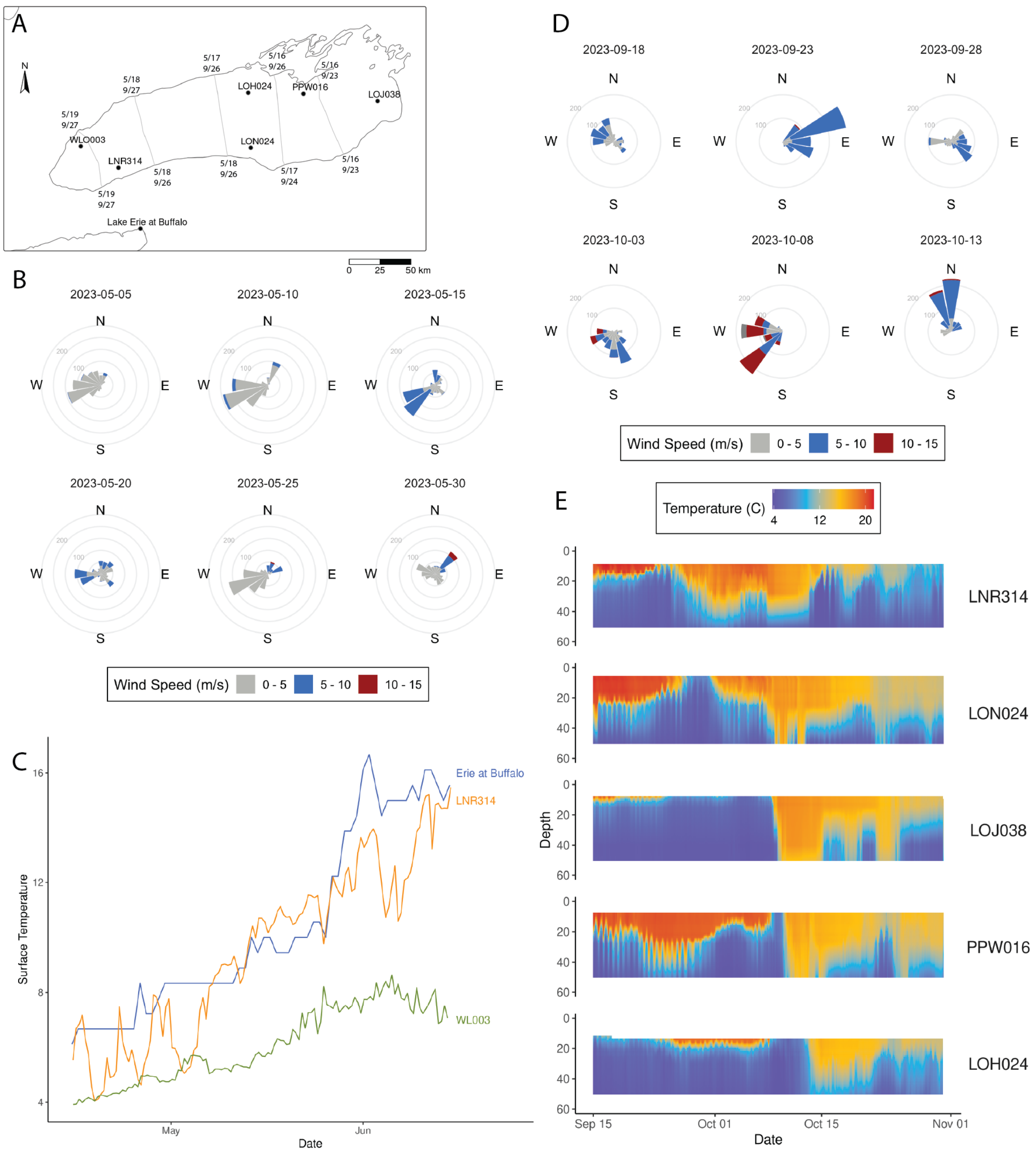


*Figure S7. Meteorological and Hydrodynamic Time Series Structuring Surface Temperature during Sampling.* (A) Map of GLATOS (in Lake Ontario) and National Weather Service temperature loggers. (B) Binned wind roses in the days leading up to and following sampling in May. (C) Time series of temperature moorings in the days leading up to and following sampling in May. The temperature sensor closest to 12 meters was selected for WLO003 and LNR314; the Lake Erie at Buffalo temperature sensor is 30m deep. (D) Binned wind roses in the days leading up to and following sampling in September. (E) Time series of temperature moorings in the days leading up to and following sampling in September. Mooring depth was capped at 50m. Temperatures were averaged in 2-hour bins. Vertical temperature profiles were linearly interpolated at each timepoint. *For wind roses:* All observations were drawn from NOAA buoy 45012. Bars represent the number of 10-minute observations of wind at a given intensity and direction. Dates above each rose correspond to the start day of a five-day period over which observations are summarized.

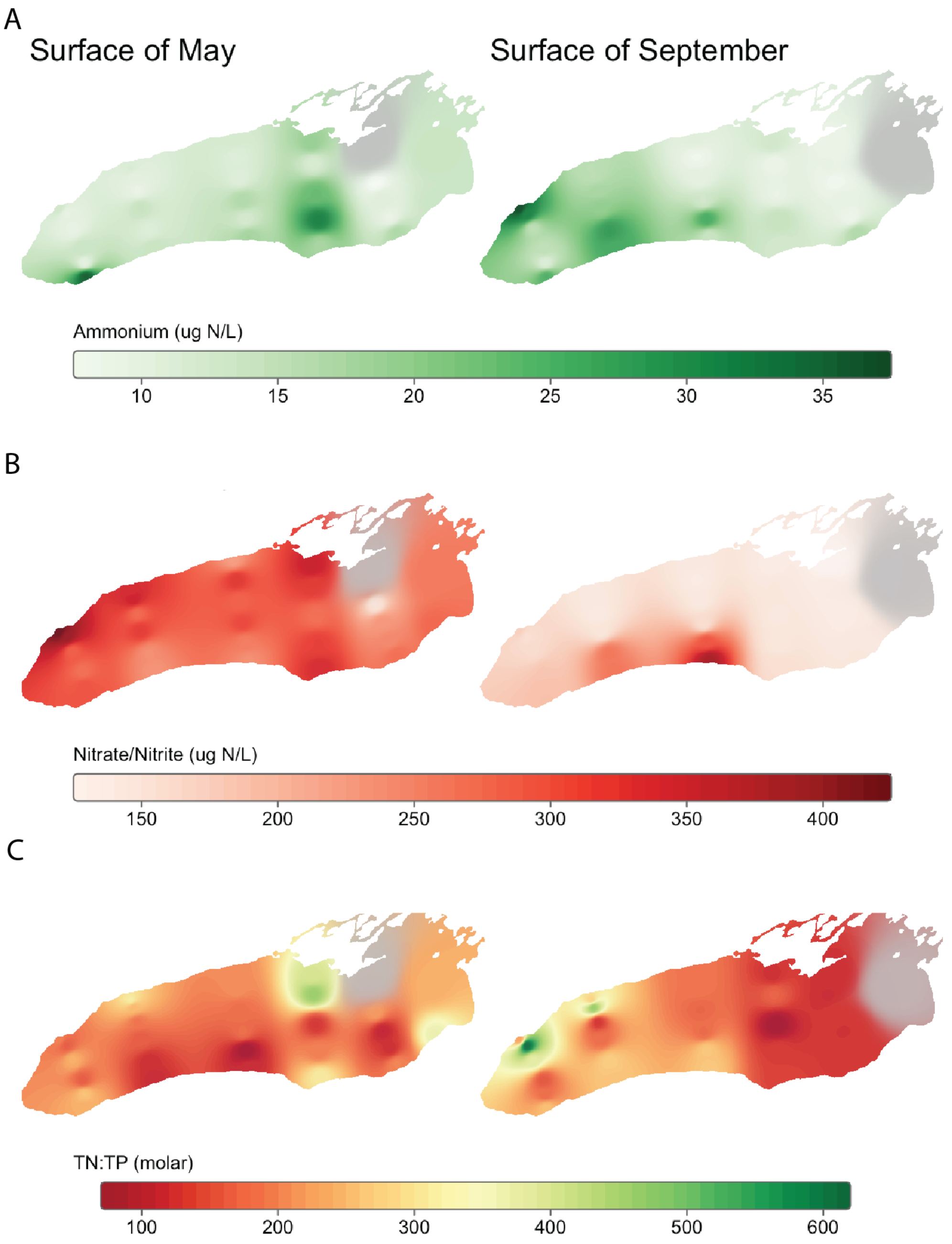


*Figure S8. Seasonal and Spatial variations of surface nitrogen species across Lake Ontario.* Multilevel B-spline interpolation of measured concentrations of (A) Ammonium ($NH_4$) (B) Nitrate and Nitrite ($NO_x$) and (C) the molar ratio of Total Nitrogen to Total Phosphorus (TN:TP) in surface samples in May and September. Interpolations were performed using inverse-distance weighted interpolation in R, with a power of 5.

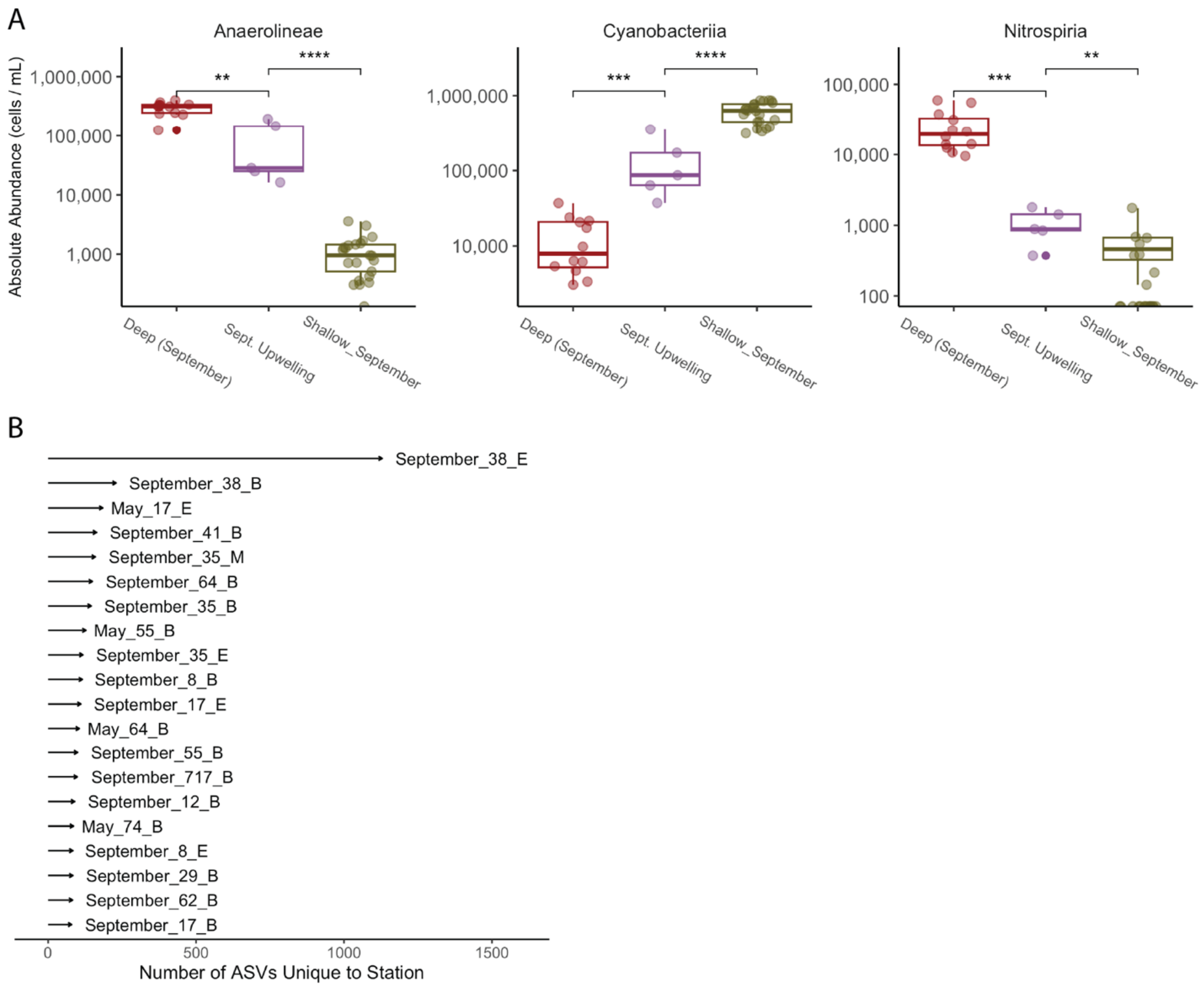


*Figure S9. Unique Taxa in the September Southern Upwelling Stations.* (A) Absolute abundances of three Classes of Bacteria in September deep, shallow, and upwelling stations. Here, stations 35 and 38 are used as September upwelling stations. Statistical tests represent results of Two-Sample Wilcoxon tests with Bonferroni correction (* = $p < 0.05$, ** = $p < 0.01$, *** = $p < 0.001$, **** = $p < 0.0001$). (B) The number of ASVs that are unique to a given sample. Any ASV which was only observed in a single sample was counted. The top twenty samples with the most unique ASVs are shown.

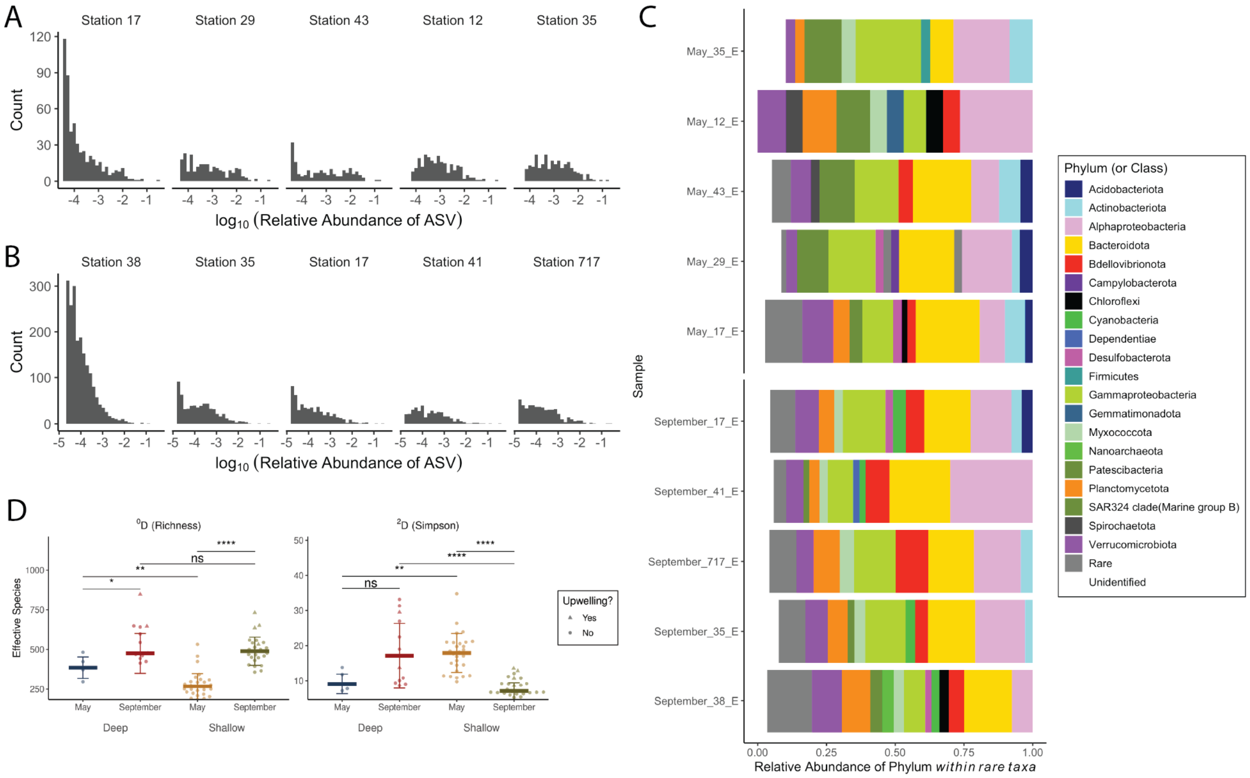


*Figure S10. Rare taxa differentiate the microbial communities of the Welland Canal and areas of Upwelling in Lake Ontario.* (A-B) Distribution of ASV abundances in anomalous stations (Welland Canal and upwelling areas) and nearby stations in (A) May and (B) September. (C) Relative abundance of rare ASVs in anomalous and nearby stations. ASVs with greater than 0.01% abundance were removed, and the relative abundance of remaining ASVs was calculated within each sample and summarized at the Phylum level. (D) Alpha diversity between depth-month groups, measured using Hill numbers within the iNEXT package. Comparisons represent Two-Sample Wilcoxon Tests (* = $p < 0.05$, ** = $p < 0.01$, *** = $p < 0.001$, **** = $p < 0.0001$).

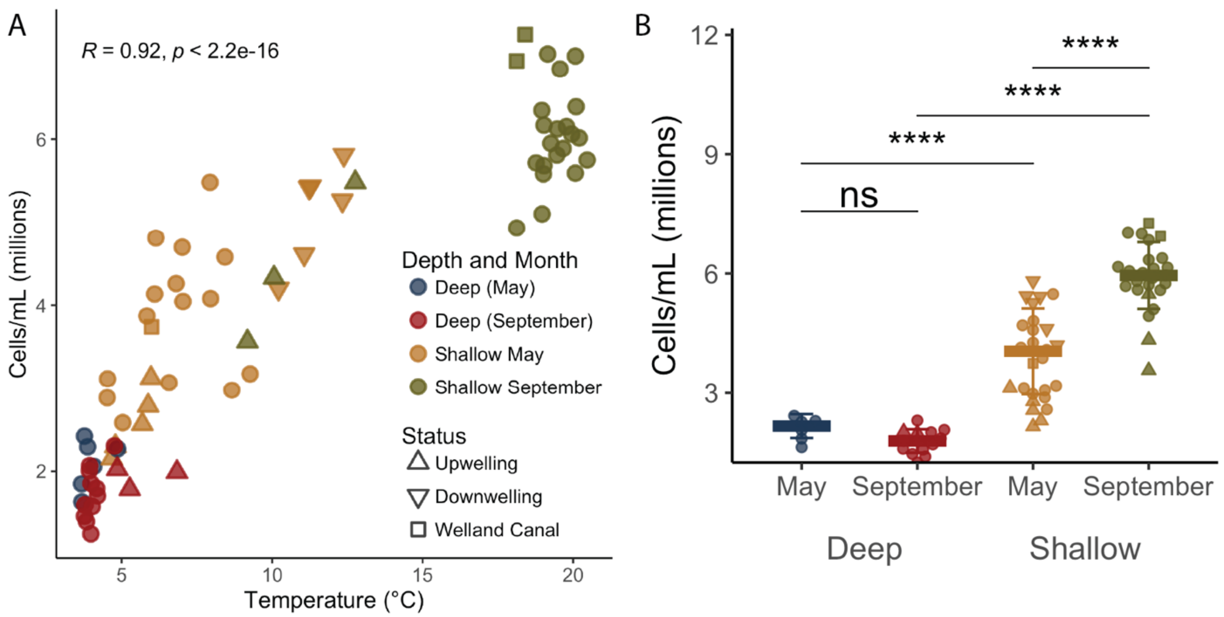


*Figure S11. Cell counts are strongly correlated with temperature.* (A) Cell abundances in millions of cells/mL (y-axis) compared to temperature in Celsius (x-axis). *R* corresponds to the Spearman correlation. (B) Cell counts split between depth-month groups. Comparisons represent Two-Sample Wilcoxon Tests (* = p < 0.05, ** = p < 0.01, *** = p < 0.001, **** = p < 0.0001).

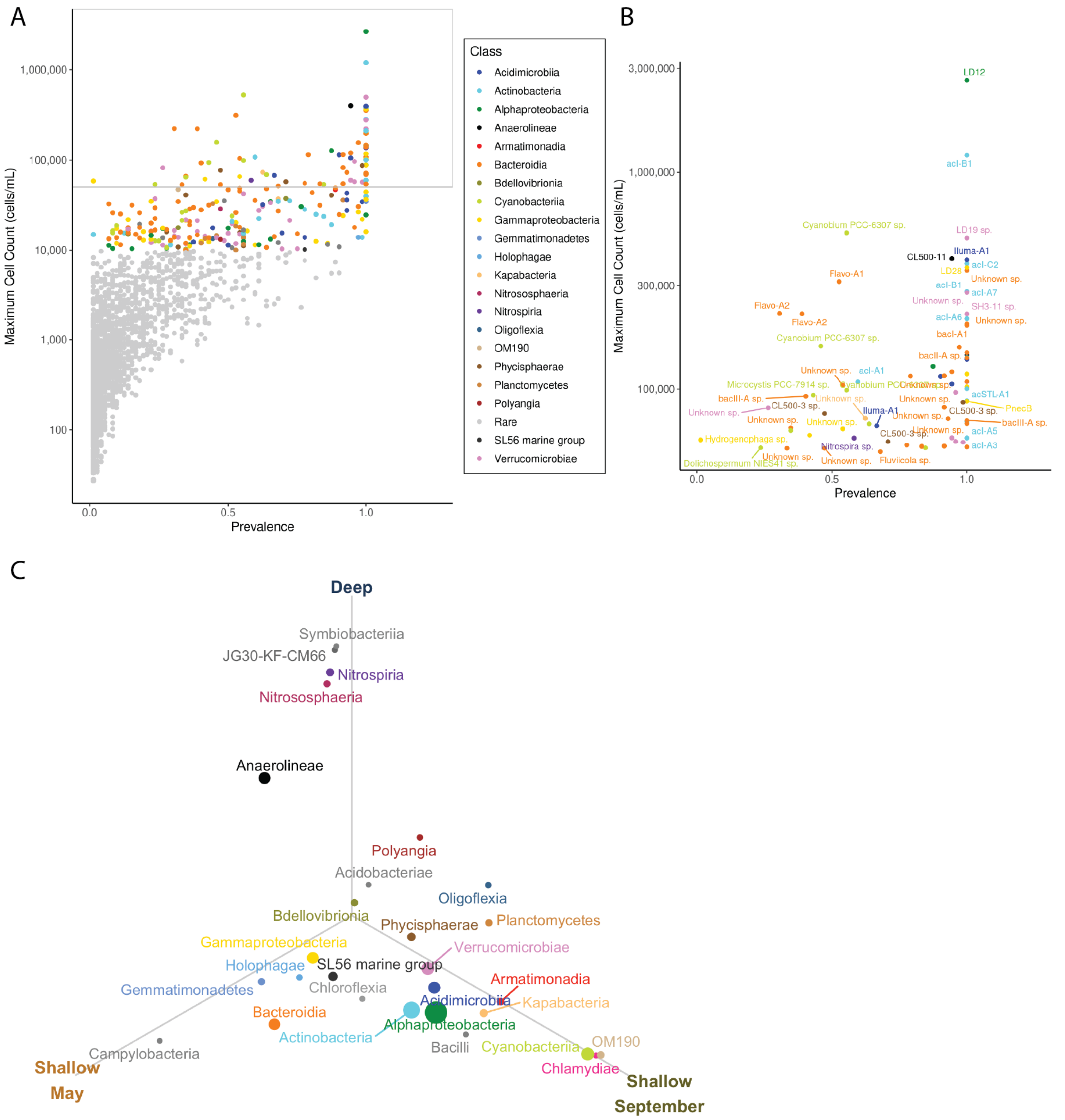


*Figure S12. Distribution and Differential Abundance of Microbial Taxa in Lake Ontario* (A) Maximum observed cell count and prevalence (percentage of samples for which that ASV was observed) for individual ASVs are plotted and colored by Class. (B) ASVs whose maximum abundance exceeds 50,000 cells/mL. Labeling reflects Genus and Species as assigned by TaxAss. (C) Differential abundance was calculated using ANCOM-BC2 using pairwise comparisons between our three month/depth groups at the Class level. For taxa which were differentially abundant ($p < 0.05$ and a passing sensitivity test) in at least one pairwise comparison were retained. A Class's location in the plot represents the centroid of a triangle whose vertices are the average absolute abundance of that class in each month/depth group.Class colors are consistent across all three plots.

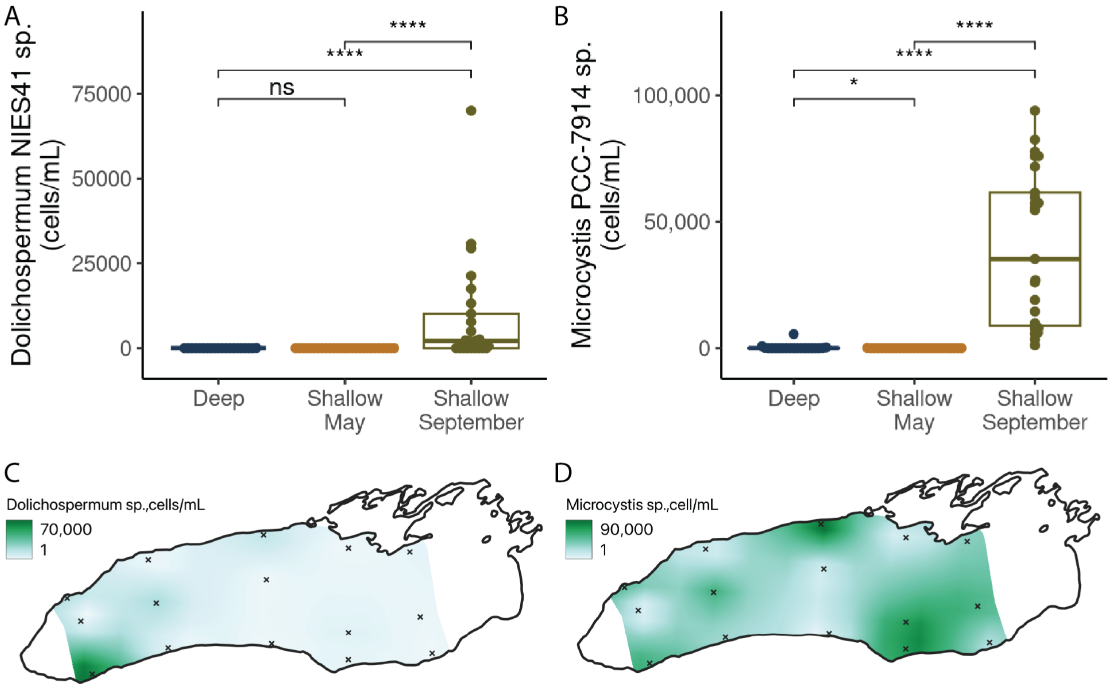


*Figure S13. Abundance of Potentially Harmful Cyanobacteria in Lake Ontario.* Absolute abundance of two genera of potentially harmful bacteria across month-depth groups, including (A) *Dolichospermum NIES41* of unknown species and (B) *Microcystis PCC-7914* of unknown species. Comparisons represent Two-Sample Wilcoxon Tests (* = $p < 0.05$, ** = $p < 0.01$, *** = $p < 0.001$, **** = $p < 0.0001$). (C-D) Spatial distribution of the (C) *Dolichospermum sp.* and (D) *Microcystis sp.* in September surface samples. Note that scales are different between both maps. Interpolation was performed using multilevel B-splines in QGIS/GRASS with 20km east-west steps, 10km north-south steps, and a Tykhonov regularization of 0.05.

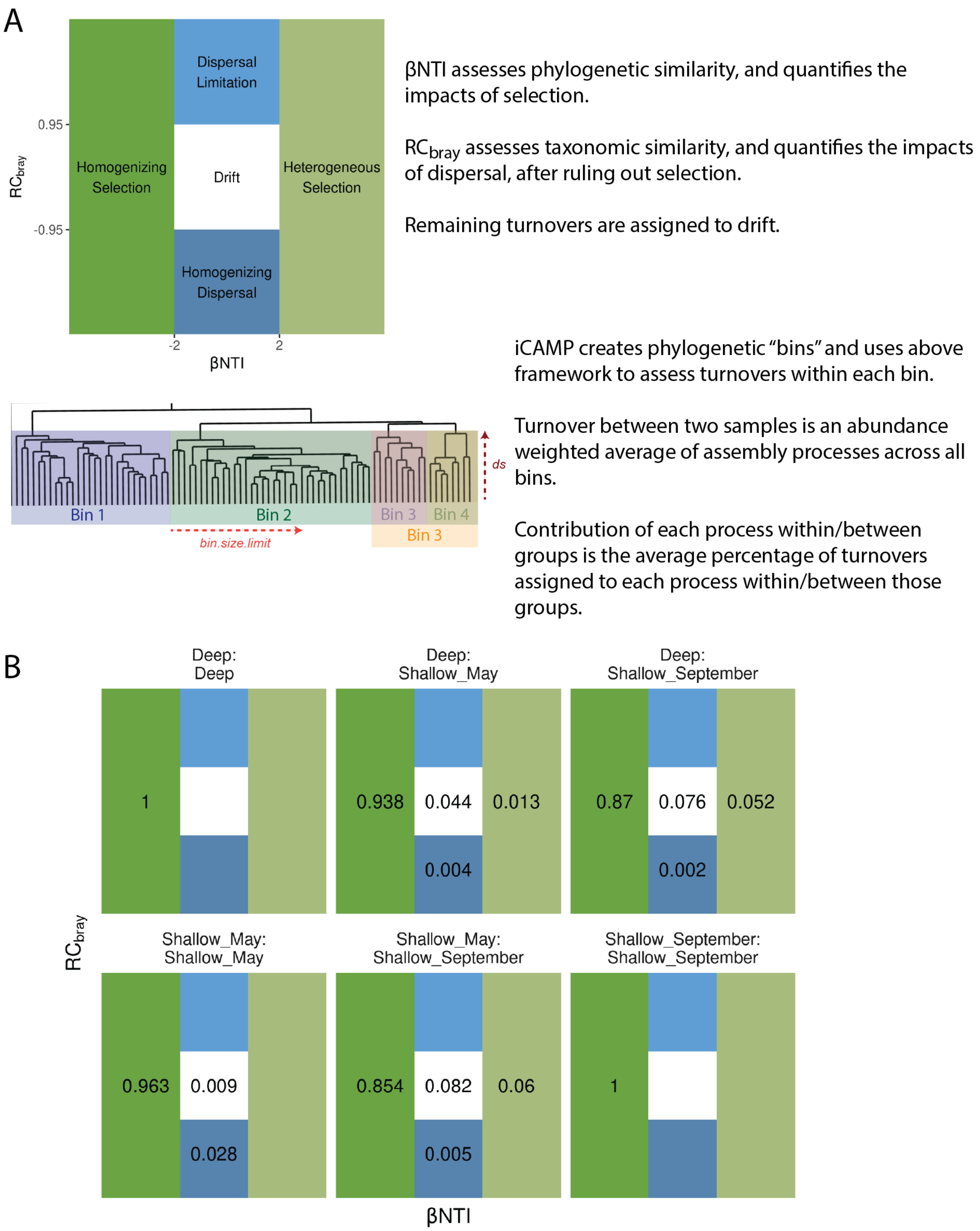


*Figure S14. Demonstration of the iCAMP framework and its comparison with classical methods for quantifying community assembly processes.* (A) iCAMP quantifies how selection, dispersal and drift shape community turnover by integrating phylogenetic (x-axis) and taxonomic (y-axis) information. It calculates β-Mean Pairwise Distance (βMPD) or β-Mean Nearest Taxon Distance (βMNTD) and their standardized forms β-Nearest Relative Index (βNRI) or β-Nearest Taxon Index (βNTI) to detect selection. When these metrics show that phylogenetic patterns are not shaped by selection, iCAMP tests for dispersal effects using the Raup-$Crick_{Bray}$ ($RC_{bray}$) index, which emphasizes taxonomic turnover. Turnovers not explained by either selection or dispersal are assigned to ecological drift. iCAMP groups ASVs into monophyletic “bins”, evaluates the dominant process for each bin based on the relative abundance of ASV represented by each bin within each sample, and then averages across bins to estimate the overall contribution of each process within or between groups. (B) Assembly processes as estimated using βNTI (($|\beta NTI| \geq 2$)) and Raup-$Crick_{Bray}$ ($|RC_{bray}| \geq 0.95$) using iCAMP::bNTIn.p and iCAMP::RC.pc functions. Numbers within each area represent the percent of pairwise turnovers assigned to each process (as illustrated in A) within or between the indicated group(s).

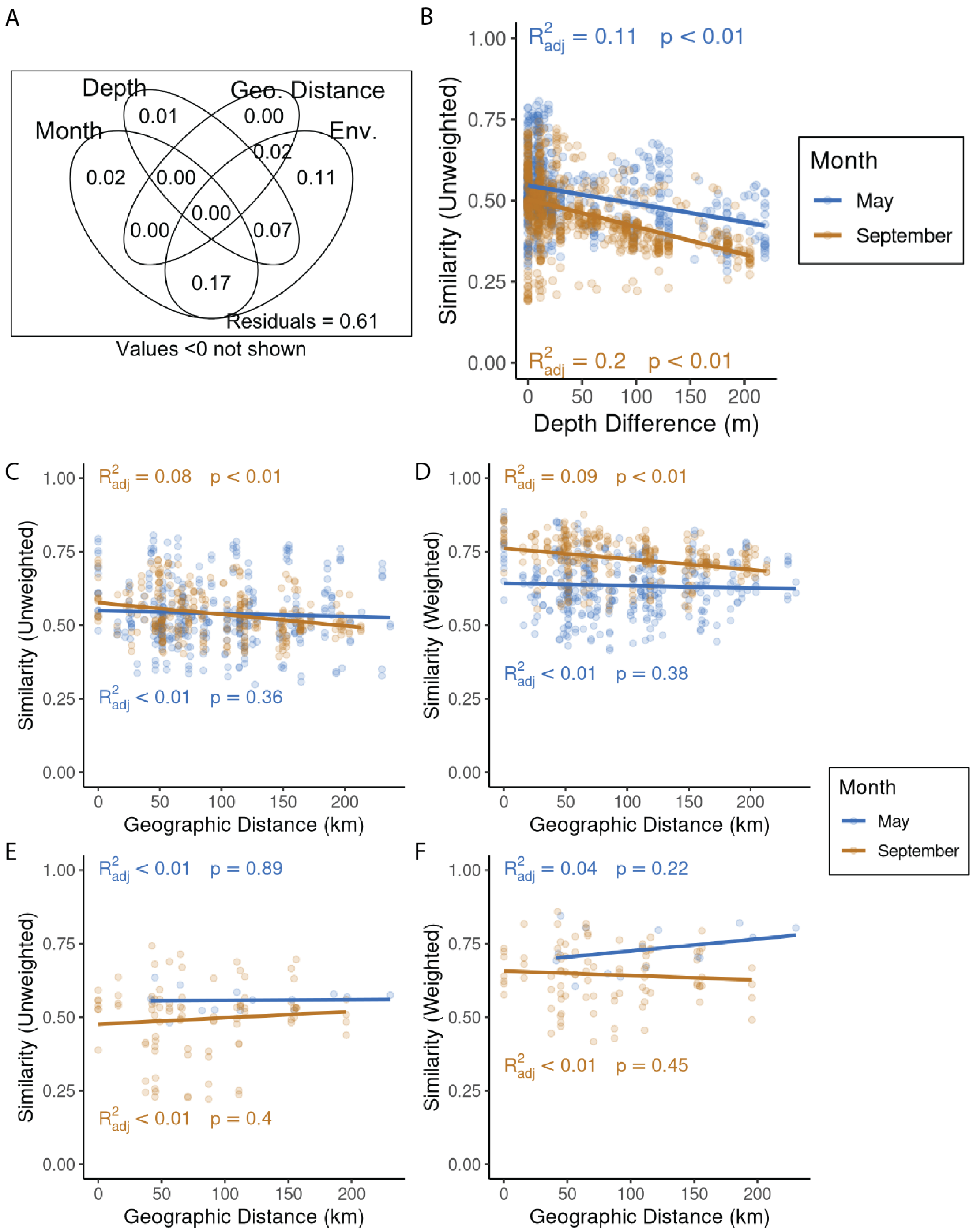


*Figure S15. Fine-grain differences in decay relationships.* (A) Variance partitioning results using the abundance-unweighted UniFrac dissimilarity as a response. Environmental variables (Env.) corresponded to scaled physical and chemical parameters from each sample. (B) Depth decay using abundance-unweighted UniFrac dissimilarity. Similarity is defined as 1 - unweighted UniFrac. Linear models were calculated separately for samples from each month. (C-F) Distance decay relationships in shallow samples (C and D) versus deep samples (E and F), and abundance-unweighted UniFrac (C and E) and abundance-weighted UniFrac (D and F).

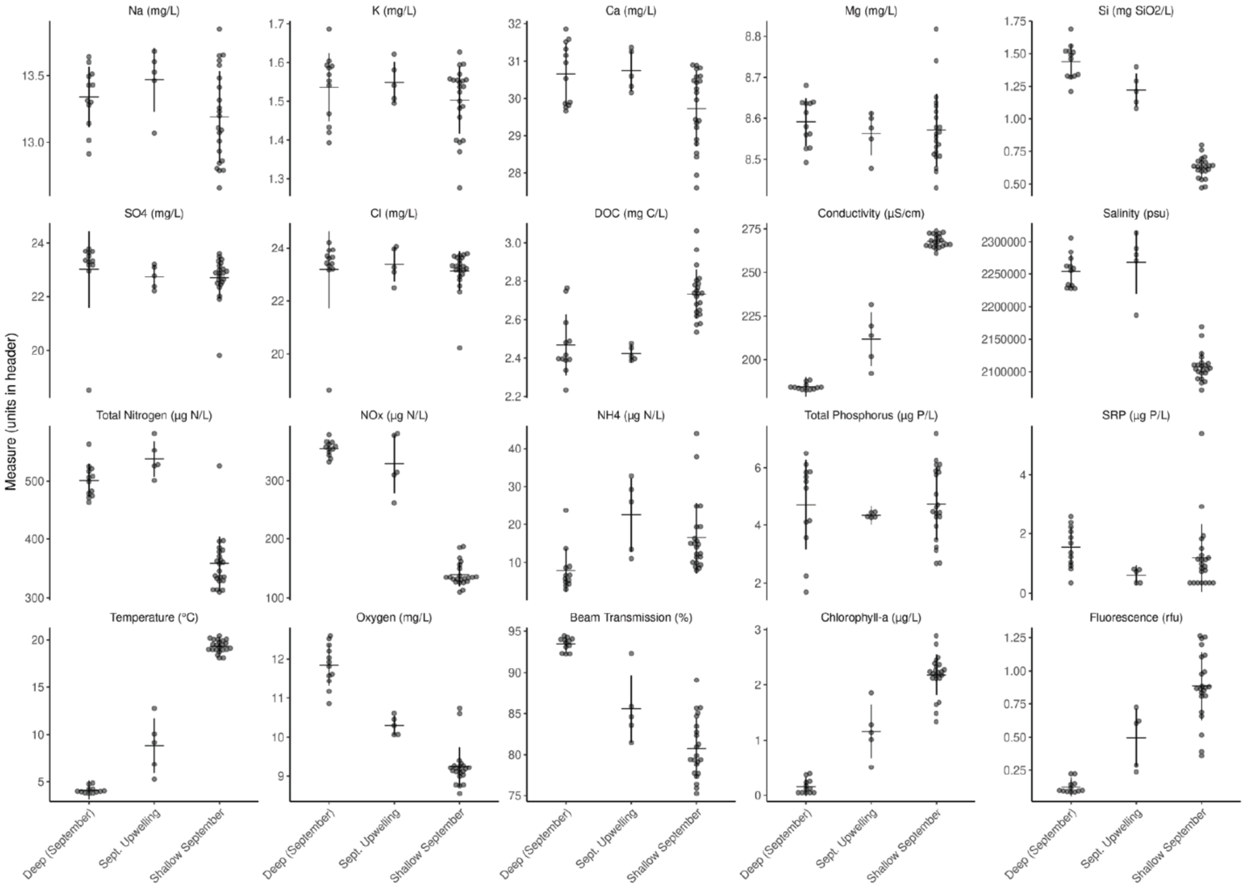


*Figure S16. Chemical and physical differences in samples from deep, shallow, and upwelling areas in September.* Each point corresponds to a sample; the center line is the mean ± the standard deviation. Sept. upwelling samples are September_38_E, September_35_E, September_38_B, September_35_M, and September_35_B.

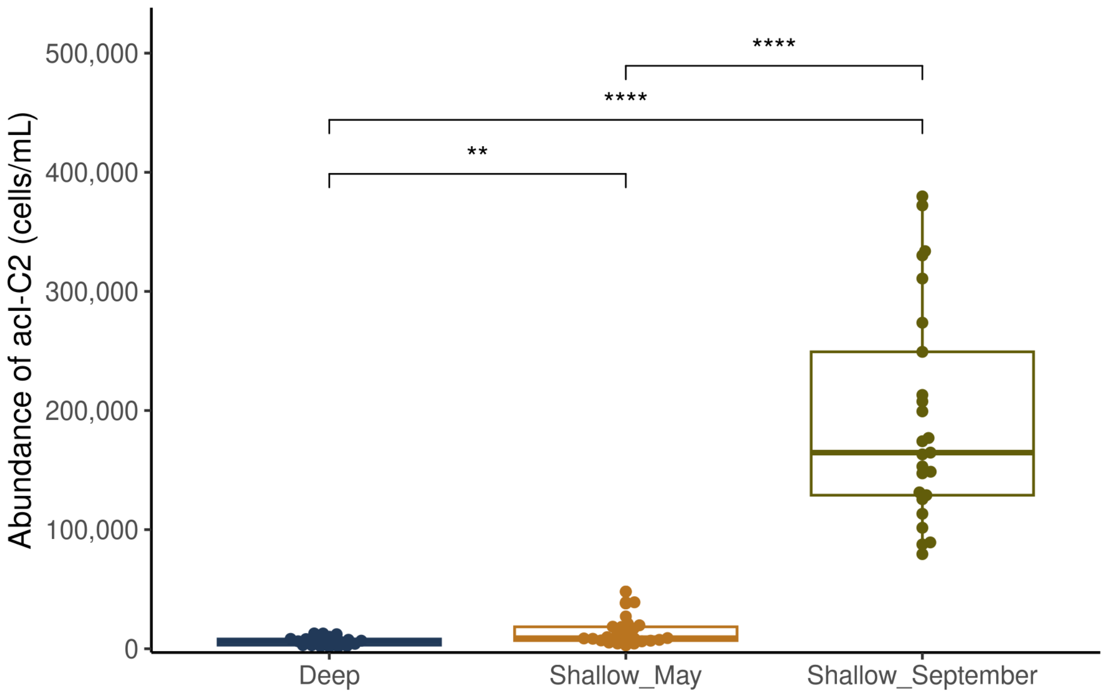


*Figure S17. Abundance of Actinobacteria acI-C2*.* Absolute abundance of Actinobacteria species *acI-C2* across three depth-month groups. Significance represents Two-sample Wilcoxon Tests with Bonferroni correction (* = $p < 0.05$, ** = $p < 0.01$, *** = $p < 0.001$, **** = $p < 0.0001$).

**Supplemental Tables**

Table S1. List of Stations for Microbial Sampling

| Station | Date | Time | Latitude | Longitude | Depths Sampled (m) | Notes |
|---|---|---|---|---|---|---|
| May Cruise | | | | | | |
| 008 | 20230519 | 1:20:00 PM | 43.6232 | -79.4518 | 5 | |
| 012 | 20230519 | 10:32:00 AM | 43.5035 | -79.3537 | 5, 15, 102.8 | |
| 017 | 20230519 | 7:10:00 AM | 43.2250 | -79.2714 | 5 | |
| 029 | 20230518 | 3:51:00 PM | 43.8184 | -78.8695 | 5, 19, 27 | |
| 033 | 20230518 | 11:35:00 AM | 43.5969 | -78.8133 | 5, 20, 135 | |
| 035 | 20230518 | 7:54:00 AM | 43.3618 | -78.7294 | 5, 10, 27 | |
| 038 | 20230518 | 2:24:00 AM | 43.3834 | -77.9896 | 5, 14 | |
| 041 | 20230517 | 8:02:00 PM | 43.7166 | -78.0273 | 5, 24, 127 | |
| 043 | 20230517 | 5:07:00 PM | 43.9501 | -78.0497 | 5 | |
| 048 | 20230516 | 5:58:00 PM | 43.8797 | -77.4397 | 5, 35 | |
| 055 | 20230517 | 10:22:00 AM | 43.4442 | -77.4389 | 5, 13, 189 | |
| 064 | 20230516 | 2:15:00 AM | 43.5197 | -76.91836 | 5, 14, 224 | |
| 066 | 20230516 | 7:25:00 AM | 43.3336 | -76.8343 | 5 | |

| Station | Date | Time | Latitude | Longitude | Depths Sampled (m) | Notes |
|---|---|---|---|---|---|---|
| 074 | 20230515 | 4:37:00 PM | 43.7473 | -76.5115 | 5, 14, 66 | |
| 717 | 20230517 | 8:00:00 AM | 43.3005 | -77.4408 | 5 | |
| September Cruise | | | | | | |
| 008 | 20230927 | 7:51:00 PM | 43.6222 | -79.4509 | 5, 13.4 | |
| 012 | 20230927 | 11:21:00 PM | 43.5034 | -79.3531 | 5, 52, 102 | |
| 017 | 20230927 | 2:54:00 PM | 43.2253 | -79.2711 | 5, 10.98 | |
| 029 | 20230927 | 4:12:00 AM | 43.8194 | -78.8711 | 5, 15, 27.9 | |
| 033 | 20230927 | 9:16:00 AM | 43.5965 | -78.8143 | 5, 68.5, 134.9 | Day Sample |
| 033 | 20230926 | 11:42:00 PM | 43.5973 | -78.8149 | 5, 68, 134 | Night Sample |
| 035 | 20230926 | 1:49:00 PM | 43.3614 | -78.7294 | 5, 14.4, 26.7 | |
| 038 | 20230926 | 10:25:00 AM | 43.3831 | -77.9887 | 5, 16.6 | |
| 041 | 20230926 | 6:39:00 AM | 43.7174 | -78.0263 | 5, 63.9, 126 | |
| 043 | 20230926 | 3:38:00 AM | 43.9489 | -78.0468 | 5, 15.6 | |
| 048 | 20230926 | 12:31:00 AM | 43.8797 | -77.4392 | 5, 16.5, 31.5 | |
| 055 | 20230925 | 7:10:00 PM | 43.4428 | -77.4398 | 5, 96, 189.6 | |

| Station | Date | Time | Latitude | Longitude | Depths Sampled (m) | Notes |
|---|---|---|---|---|---|---|
| 062 | 20230923 | 3:11:00 PM | 43.5251 | -76.9265 | 5, 18.3 | |
| 064 | 20230923 | 11:50:00 AM | 43.3332 | -76.8413 | 5, 106, 210 | |
| 066 | 20230923 | 9:19:00 PM | 43.8600 | -77.0012 | 5, 15.8 | |
| 717 | 20230924 | 9:55:00 AM | 43.3004 | -77.4407 | 5, 18.5, 34.8 | |

393

Table S2. Software Versions and Citations

| Software | Version | Citation |
|---|---|---|
| **Languages and Environments** | | |
| R | 4.3.2 | [31] |
| RStudio | 2022.07.0 | [32] |
| QGIS | 3.40.5 | [33] |
| **CLI Tools** | | |
| MAFFT | 7.520 | [34] |
| TaxAss | 2.1.1 | [3] |
| FAPROTAX | 1.2.10 | [35] |
| **R Packages** | | |
| ape | 5.7-1 | [36] |
| ANCOMBC | 1.6.4 | [37] |
| biomformat | 1.30.0 | [38] |
| Biostrings | 2.64.1 | [39] |
| broom | 1.0.5 | [40] |
| combinat | 0.0-8 | [41] |
| dada2 | 1.24.0 | [2] |
| flowCore | 2.14.2 | [42] |
| ggcyto | 1.30.2 | [43] |
| ggdendro | 0.2.0 | [44] |
| ggfortify | 0.4.16 | [45] |
| ggh4x | 0.2.8 | [46] |
| ggpubr | 0.6.0 | [47] |
| ggrepel | 0.9.4 | [48] |
| ggside | 0.3.1 | [49] |
| ggtree | 3.4.4 | [50] |

| Software | Version | Citation |
|---|---|---|
| gridExtra | 2.3 | [51] |
| gstat | 2.1-1 | [52] |
| iCAMP | 1.5.12 | [53] |
| iNEXT | 3.0.0 | [54] |
| lubridate | 1.9.3 | [55] |
| microViz | 0.12.0 | [56] |
| MBA | 0.1-0 | [57] |
| oce | 1.8-2 | [58] |
| pacman | 0.5.1 | [59] |
| patchwork | 1.2.0 | [60] |
| phyloseq | 1.40.0 | [61] |
| phytools | 2.1-1 | [62] |
| purrr | 1.0.2 | [63] |
| readxl | 1.4.3 | [64] |
| rstatix | 0.7.2 | [65] |
| scales | 1.3.0 | [66] |
| sf | 1.0-14 | [67] |
| speedyseq | 0.5.3.9018 | [68] |
| terra | 1.7-71 | [69] |
| tidytree | 0.4.6 | [70] |
| tidyverse | 2.0.0 | [71] |
| tmap | 3.99.9 | [72] |
| tmaptools | 3.1-1 | [73] |
| treemapify | 2.5.6 | [74] |
| units | 0.8-5 | [75] |
| vegan | 2.6-4 | [76] |

31. R Core Team. R: A Language and Environment for Statistical Computing. Vienna, Austria: R Foundation for Statistical Computing, 2020.

32. RStudio Team. RStudio: Integrated Development Environment for R. Boston, MA: RStudio, PBC., 2020.

33. QGIS Development Team. QGIS Geographic Information System. 2021. 2021.

34. Katoh K, Standley DM. MAFFT Multiple Sequence Alignment Software Version 7: Improvements in Performance and Usability. *Molecular Biology and Evolution* 2013;**30**:772–780. https://doi.org/10.1093/molbev/mst010

35. Louca S, Parfrey LW, Doebeli M. Decoupling function and taxonomy in the global ocean microbiome. *Science* 2016;**353**:1272–1277. https://doi.org/10.1126/science.aaf4507

36. Paradis E, Schliep K. ape 5.0: an environment for modern phylogenetics and evolutionary analyses in R. *Bioinformatics* 2019;**35**:526–528. https://doi.org/10.1093/bioinformatics/bty633

37. Lin H. ANCOMBC: Microbiome differential abudance and correlation analyses with bias correction. 2022.

38. McMurdie PJ, Paulson JN. biomformat: An interface package for the BIOM file format. 2023.

39. Pagès H et al. Biostrings: Efficient manipulation of biological strings. 2022.

40. Robinson D, Hayes A, Couch S. broom: Convert statistical objects into tidy tibbles. 2023.

41. Chasalow S. combinat: combinatorics utilities. 2012.

42. Ellis B et al. flowCore: flowCore: Basic structures for flow cytometry data. 2024.

43. Jiang M. ggcyto: Visualize Cytometry data with ggplot. 2024.

44. de Vries A, Ripley BD. ggdendro: Create dendrograms and tree diagrams using ggplot2. 2024.

45. Horikoshi M, Tang Y. ggfortify: Data visualization tools for statistical analysis results. 2023.
46. van den Brand T. ggh4x: Hacks for ‘ggplot2’. 2024.
47. Kassambara A. ggpubr: ggplot2 based publication ready plots. 2023.
48. Slowikowski K. ggrepel: Automatically Position Non-Overlapping Text Labels with ‘ggplot2’. 2022.
49. Landis J. ggside: Side grammar graphics. 2024.
50. Xu S et al. Ggtree: A serialized data object for visualization of a phylogenetic tree and annotation data. *iMeta* 2022;**1**:e56. https://doi.org/10.1002/imt2.56
51. Auguie B. gridExtra: Miscellaneous functions for ‘Grid’ graphics. 2017.
52. Gräler B, Pebesma E, Heuvelink G. Spatio-temporal interpolation using gstat. *The R Journal* 2016;**8**:204–218.
53. Ning D. iCAMP: Infer community assembly mechanisms by phylogenetic-bin-based null model analysis. 2022.
54. Hsieh TC, Ma KH, Chao A. iNEXT: Interpolation and extrapolation for species diversity. 2022.
55. Spinu V, Grolemund G, Wickham H. lubridate: Make dealing with dates a little easier. 2023.
56. Barnett D. microViz: Microbiome data analysis and visualization. 2024.
57. Finley A, Banerjee S, Hjelle Ø. MBA: Multilevel b-spline approximation. 2022.
58. Kelley D, Richards C. oce: Analysis of oceanographic data. 2023.
59. Rinker TW, Kurkiewicz D. pacman: Package management for R. Buffalo, New York, 2018.
60. Pedersen TL. patchwork: The composer of plots. 2024.

61. McMurdie PJ, Holmes S. phyloseq: An R package for reproducible interactive analysis and graphics of microbiome census data. *PLoS ONE* 2013;**8**:e61217.
62. Revell LJ. phytools: An R package for phylogenetic comparative biology (and other things). *Methods in Ecology and Evolution* 2012;**3**:217–223.
63. Wickham H, Henry L. purrr: Functional programming tools. 2023.
64. Wickham H, Bryan J. readxl: Read excel files. 2023.
65. Kassambara A. rstatix: Pipe-friendly framework for basic statistical tests. 2023.
66. Wickham H, Pedersen TL, Seidel D. scales: Scale functions for visualization. 2023.
67. Pebesma E. sf: Simple features for R. 2023.
68. McLaren M. speedyseq: Faster implementations of phyloseq functions. 2024.
69. Hijmans RJ. terra: Spatial data analysis. 2024.
70. Yu G. tidytree: A tidy tool for phylogenetic tree data manipulation. 2023.
71. Wickham H. tidyverse: Easily install and load the tidyverse. 2023.
72. Tennekes M. tmap: Thematic maps. 2024.
73. Tennekes M. tmaptools: Thematic map tools. 2021.
74. Wilkins D. treemapify: Draw treemaps in ‘ggplot2’. 2023.
75. Pebesma E, Mailund T, Hiebert J. Measurement units in R. *R Journal* 2016;**8**:486–494. https://doi.org/10.32614/RJ-2016-061
76. Oksanen J et al. vegan: Community ecology package. 2022.
